# Systematic De-Risking of TCR-Mimic Therapeutics Through Proteome-Wide Off-Target Landscaping and a Generalizable Design Rule Framework

**DOI:** 10.64898/2026.07.30.741432

**Authors:** Simon Schuster, Frederike A. Hartl, Jennifer Stehle, Fabian Beier, Hoor Al-Hasani, Christoph Reinhart, Tsai-Hsuan Weng, Stefan Krämer, Pranav Hamde, Tobias Herz, Sebastian Österlin, Marco Schmidt, Tabea Gross, Luca Stehl, Normann Kilb, Gloria Jünemann, Kirsten Heiss, Alexander Sinclair, Juliane Schoettgen, Philipp Meyer, Bilge Atay, Badeel Kh Q Zaghla, Steve Link, Julian Goll, Benedetta Leoni, Björn Steinmann, Oliver Selinger, Sonja Welsch, Guenter Roth, Hartmut Michel, Martin G. Klatt, Joerg Birkenfeld

## Abstract

TCR-mimic (TCRm) antibodies targeting peptide-human leukocyte antigen (pHLA) complexes enable precision immunotherapy against intracellular antigens, including cancer-testis antigens (CTAs). Achieving high specificity, however, remains challenging because of the vast diversity of the human immunopeptidome and the associated risk of off-target recognition. Here, we introduce ValidaTe, a unified framework for the proteome-scale prediction, validation, and mitigation of off-target liabilities in pHLA-directed therapeutics. ValidaTe integrates rational target prioritization, peptide-centric binder selection, proteome-wide off-target prediction, and therapeutic engineering into a hierarchical de-risking workflow. Using the CTA MAGE-A4 as a proof-of-concept, we identify the TCRm antibodies VR-4 and VR-6 with superior specificity and demonstrate how this workflow enables the discovery of safer pHLA-targeted binders. Furthermore, ValidaTe establishes the basis for the WiFi (Widened Fingerprint) engineering principle, which rationally combines TCRms with complementary off-target fingerprints in trivalent T-cell engagers to minimize unintended interactions while preserving potent target-specific activity. Together, these findings establish a generalizable framework for the rational development of safer and more selective pHLA-targeted therapeutics. We further discuss how orthogonal proteomic characterization may complement this workflow as a final layer of translational safety assessment prior to clinical development.

**Teaser:** ValidaTe accelerates safe pHLA-targeted immunotherapy through proteome-wide off-target mapping and WiFi design

## INTRODUCTION

Most intracellular proteins are continually processed by the proteasome into short peptide fragments that are loaded onto human leukocyte antigen (HLA) class I molecules to form peptide-HLA class I (pHLA-I) complexes on the cell surface. These complexes, composed of an HLA-I heavy chain, β2-microglobulin, and an 8 to12-amino-acid peptide, constitute the HLA-I immunopeptidome, which provides a real-time molecular readout of the intracellular proteome [1]. The term immunopeptidome can also refer to peptides presented by HLA class II molecules, but the present work focuses specifically on the HLA-I immunopeptidome, the principal substrate for CD8+ T-cell surveillance. Cytotoxic CD8+ T-lymphocytes interrogate this landscape through clonotypic T-cell receptors (TCRs) to detect infection or malignant transformation.

The exquisite specificity of T-cell receptor (TCR) recognition of peptide-human leukocyte antigen (pHLA) complexes enables precise discrimination between self and non-self antigens, including oncogenic neoepitopes and cancer-testis antigens (CTAs) that are aberrantly expressed in malignant tissues while being largely absent from healthy cells [1–4]. Consequently, the tumor HLA class I immunopeptidome provides access to a broad repertoire of intracellular targets that remain inaccessible to conventional antibody-based therapeutics or chimeric antigen receptor (CAR) modalities, which are generally limited to cell-surface antigens [5]. Therapeutic approaches exploiting tumor-specific pHLA complexes, including TCR-engineered T-cells (TCR-T), soluble TCR-based bispecific T-cell engagers, and TCR-mimic (TCRm)-based modalities such as bispecific antibodies and next-generation CAR-T platforms, are redefining translational oncology by enabling selective targeting of intracellular antigens in patients with otherwise limited therapeutic options [6, 7].

Tebentafusp, a gp100 specific engineered soluble TCR fused to an anti-CD3 recruitment domain, has demonstrated a clinically significant overall survival benefit in metastatic uveal melanoma and established proof of principle for pHLA directed therapy in solid tumors [8, 9]. More broadly, cancer testis antigen derived pHLA complexes have emerged as attractive therapeutic targets for engineered TCRs and TCR mimic modalities [4]. Supporting the clinical feasibility of this approach, the MAGE-A4 cancer testis antigen [10] has emerged as a particularly compelling target, exemplified by the FDA approval of the MAGE-A4 specific TCR T-cell therapy afamitresgene autoleucel for synovial sarcoma, further validating cancer testis antigens as clinically relevant targets for pHLA directed immunotherapies [11].

Despite these advances, a fundamental challenge persists. Most tumor-associated pHLA-I complexes are presented at low copy numbers, necessitating high-affinity targeting domains. Improving affinity, however, increases the risk of unintended recognition of structurally related self-peptides [12]. Clinical experience underscores the seriousness of this liability. Affinity-enhanced TCRs specific for the MAGE-A3-derived peptide EVDPIGHLY presented by HLA-A01:01 caused fatal cardiac toxicity due to cross-recognition of a Titin-derived peptide [13]. Neurologic toxicity in a separate clinical setting was attributed to recognition of peptides structurally related to the MAGE-A3-derived peptide KVAELVHFL presented by HLA-A*02:01, most likely including peptides derived from MAGE-A12 and ESPL8 [14]. These cases highlight the need for comprehensive and mechanistically grounded off-target assessment for both engineered TCRs and TCRm molecules.

Numerous computational and experimental strategies have been developed to predict off target pHLA recognition, including sequence similarity searches based on Xs-canning substitution profiles, high throughput pHLA library screening, genetic screening approaches, and structural or bioinformatics-based prediction pipelines [6, 15–17]. However, none of these methods provides sufficient breadth, resolution, and biological relevance to capture the full spectrum of pHLA class I complexes that may be recognized by engineered TCRs or TCR mimic antibodies. We recently demonstrated that machine learning models incorporating kinetic TCRm-pHLA interaction landscapes improve the detection of sequence unrelated cross reactivities [18]. Nevertheless, a unified framework that integrates computational prediction with systematic experimental validation is still lacking.

Here, we introduce ValidaTe, a unified platform for proteome scale prediction and validation of off target liabilities for pHLA directed therapeutics, with an emphasis on TCR mimic antibodies. TCRm molecules offer several advantageous biopharmaceutical properties, including compatibility with established antibody discovery, engineering, and manufacturing platforms, as well as predictable developability profiles that facilitate industrial scale optimization [19]. ValidaTe integrates machine learning guided prediction of off target pHLA interactions, high throughput kinetic profiling of candidate off targets, and proteome wide off target identification based on TCRm pHLA kinetic interaction landscapes. These complementary datasets are further integrated with structural analyses to generate comprehensive off target liability maps for individual TCRms. Using this platform, we identified the MAGE-A4-specific TCRms VR-4 and VR-6 as lead candidates with superior target specificity.

ValidaTe also establishes the conceptual basis for a new engineering principle we term WiFi (Widened Fingerprint). WiFi enables the rational combination of TCRm molecules with complementary off-target fingerprints to minimize unintended interactions while maintaining high-affinity recognition of the intended pHLA target. Using anti-MAGE-A4 WiFi T-cell engagers as an example, we demonstrate improved selectivity and a markedly reduced off-target signature compared with existing TCRm candidates, highlighting the potential of this framework to accelerate development of safer and more effective pHLA-targeted therapeutics.

## RESULTS

### Introduction to ValidaTe

To address the need for proteome-wide off-target risk assessment in TCR-like therapeutics and to systematically de-risk their pre-clinical and clinical development, we developed ValidaTe. ValidaTe is an integrated, platformized workflow combining high-throughput kinetic profiling of TCRm-pHLA interactions, ML-based proteome-wide off-target prediction, AI-driven structural modeling, and deep functional validation in cell-based translational assays (Figure 1). ValidaTe comprises two complementary ML frameworks: (i) a target peptide-centric off-target prediction model (EpiTox) [20] and (ii) a TCRm antibody-specific, proteome-wide off-target prediction model trained on high-throughput kinetic binding data (EpiPredict) [18] generated by high-density pHLA microarray screening (HighSCORE) [21]. Predicted off-target candidates are further evaluated through an AI-driven structural modeling pipeline, which assesses the structural plausibility of TCRm-pHLA complexes to refine and prioritize hits prior to experimental follow-up. This computational framework is embedded within a pHLA-centric modeling pipeline (ParaPredict) and integrated with extensive experimental validation workflows, enabling iterative refinement of both predictive performance and functional assessment. Importantly, ValidaTe is designed to identify potential cross-reactive peptides across the full human proteome, including sequences with minimal or no primary sequence homology to the intended target, a capability that addresses a critical blind spot of conventional similarity-based screening approaches and provides a systematic strategy to de-risk the development of pHLA-directed TCR-like therapeutics.

**Figure 1.**
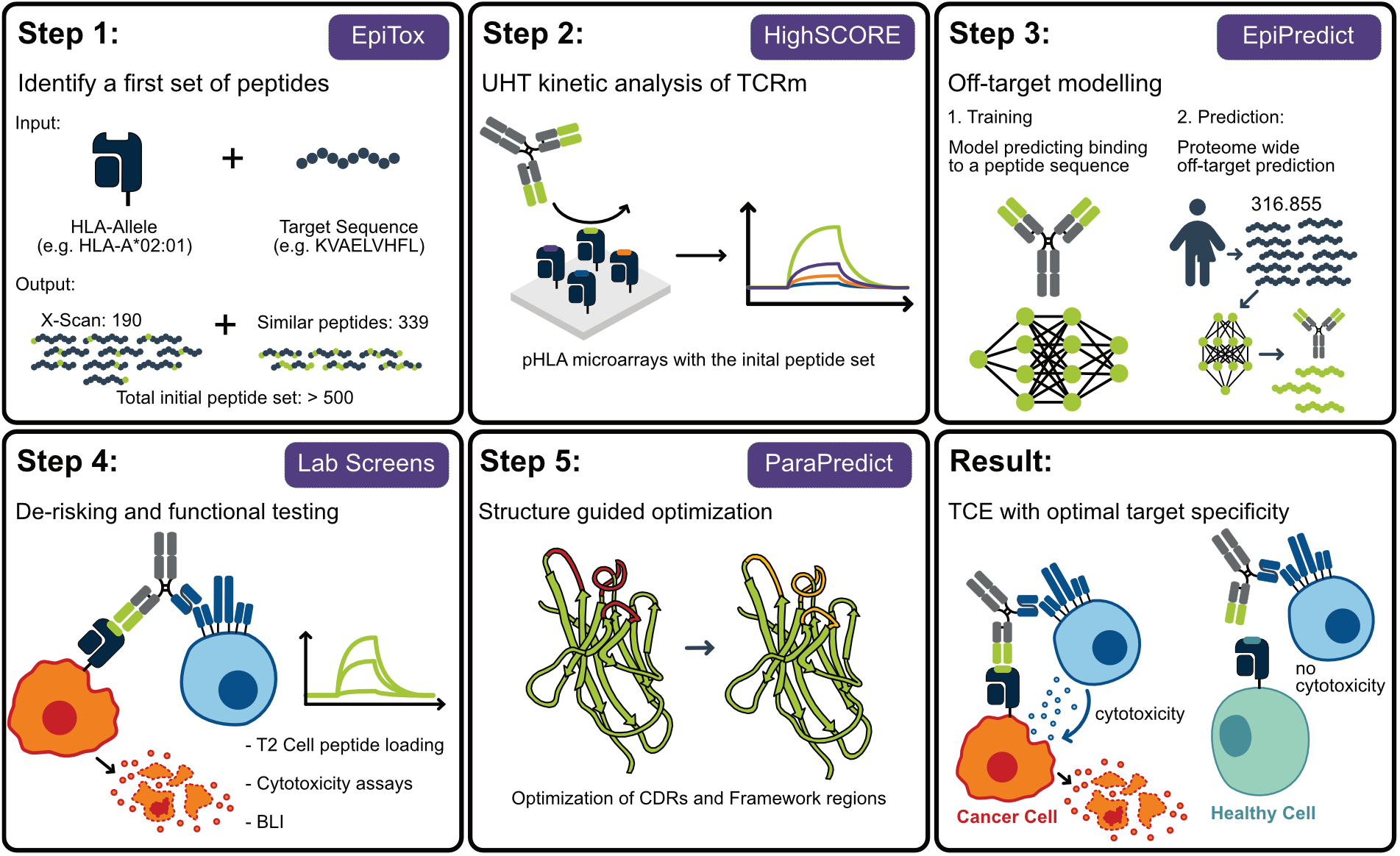
The ValidaTe Platform. ValidaTe integrates five sequential steps to identify de-risked TCRm-based T-cell engagers. EpiTox (Step 1) performs peptide-centric off-target toxicity prediction to define a physiologically relevant off-target peptide set. HighSCORE (Step 2) maps kinetic pHLA-TCRm interactions across this candidate peptide panel. EpiPredict (Step 3) applies ML-based, TCRm-centric, proteome-wide off-target prediction, inferred from HighSCORE kinetic landscapes, to prioritize TCRm leads. Top candidates are reformatted into bivalent and trivalent T-cell engagers and evaluated in T-cell-mediated cytotoxicity assays (Step 4). ParaPredict (Step 5) applies AI-driven structural modeling for structure-guided TCRm-pHLA complex analysis and TCRm optimization to enhance therapeutic performance.

In the following sections, we describe how ValidaTe and its components were applied to identify anti-MAGE-A4 TCRms with improved safety and efficacy profiles.

### Systematic Identification of Optimal MAGE-A4 pHLA Targets

An optimal peptide-HLA target for TCR-mimic (TCRm) antibody generation should combine a structurally accessible, peptide-centric epitope surface that enables highly specific molecular recognition, a favorable off-target landscape with minimal physiologically relevant sequence- or biophysically similar peptides, and robust evidence of tumor-associated presentation, ideally supported by immunopeptidomics across multiple cancer types.

The cancer-testis antigen (CTA) MAGE-A4 is broadly expressed across diverse tumor entities while being largely absent from healthy tissues [10]. Two HLA-A*02:01-presented MAGE-A4-derived peptides, KVLEHVVRV (amino acids 286-294; KVL) and GVYDGREHTV (amino acids 230-239; GVY), are currently under clinical evaluation. KVL is targeted in phase I studies using TCR-based T-cell engagers (registered as NCT05359445 [22]), whereas GVY is preclinically pursued via CAR-T [23] and clinically via a TCRm-based T-cell engager (registered as NCT06402201 [24]). We therefore evaluated both epitopes for their structural and functional suitability as TCRm targets.

#### Off-target landscape mapping using EpiTox profiling

A critical requirement for TCR-like therapeutics is minimization of off-target peptide recognition. To systematically characterize the landscape of physiologically relevant off-targets for KVL and GVY, we applied EpiTox, our machine learning-driven multimodal framework for peptide-centric off-target prediction [20]. EpiTox identifies potentially cross-reactive peptides through proteome-wide similarity searches, integrating experimentally derived and computationally predicted parameters, including tissue expression profiles, immunopeptidomics evidence of pHLA-I presentation, predicted HLA-A*02:01 binding affinity, and biophysical peptide surface similarity metrics. The framework generates prioritized sets of physiologically relevant candidate off-target peptides for preclinical safety assessment of TCR-like therapeutics. As shown in Figure 2 A, a considerably greater proportion of KVL off-targets than GVY off-targets scored above threshold on both the Bi-feature and Multi-feature axes (23.3% vs. 9.39% proportion test: *p* =*<* 0.001), indicating that KVL’s off-target pool contains substantially more peptides that are simultaneously high-affinity binders and sequence- and physiochemically similar to the target. Among these high-scoring peptides, the proportion further classified into the risk core (Broad and Critical or Critical only expression tiers) did not differ significantly between targets (47.4% vs. 47.6%, proportion test: *p* = 1), indicating that expression-based tissue filtering is applied comparably to both targets’ quadrant peptides (see Material and Methods). Consequently, the larger risk core observed for KVL reflects a larger pool of high-scoring candidates entering the expression filter, rather than a differential effect of expression criteria between targets. Together, these results show that GVY displays a markedly more restricted physiologically relevant off-target landscape than KVL, consistent with a more favorable profile for preclinical safety assessment and TCRm development.

**Figure 2.**
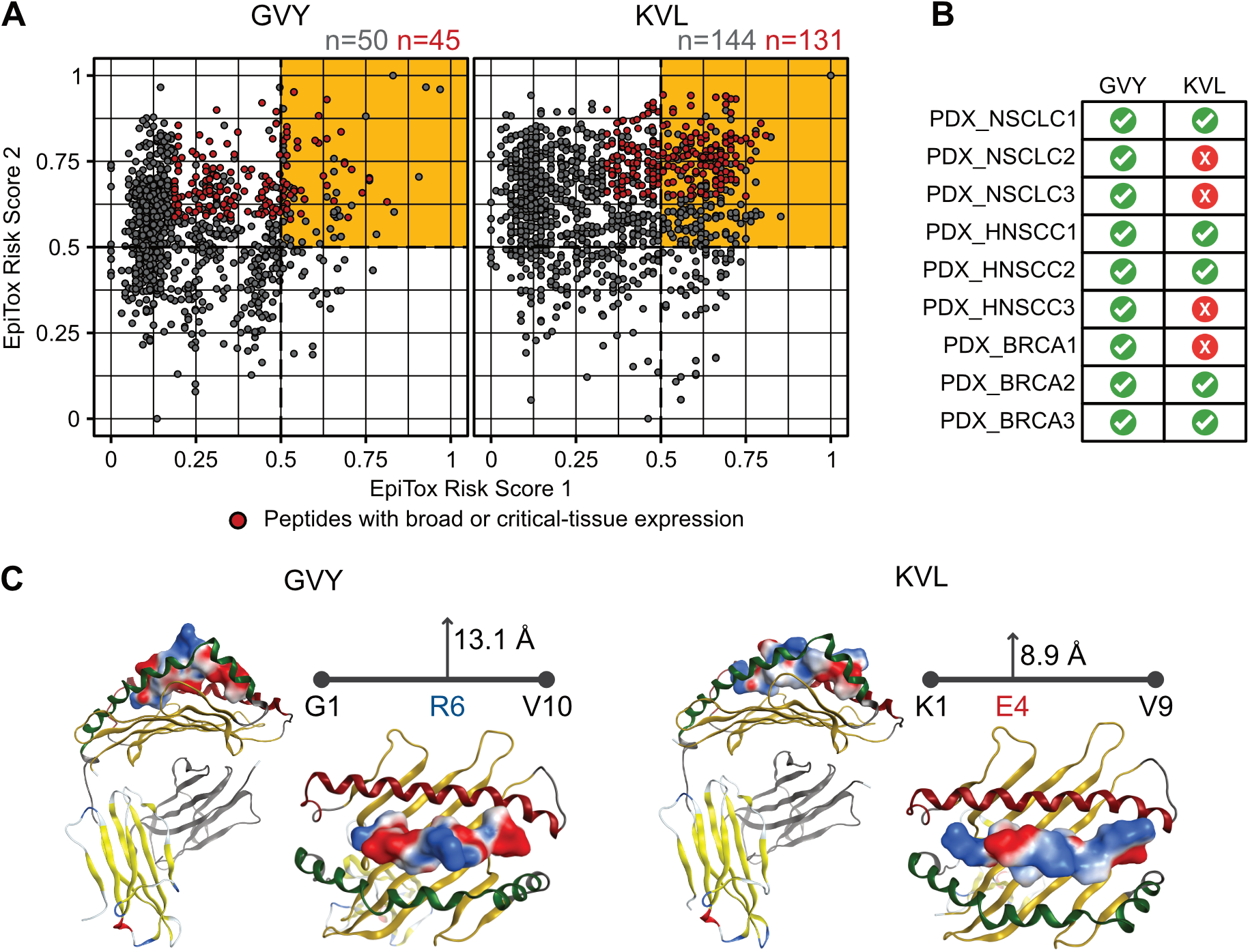
Integrated identification of optimal MAGE-A4 pHLA targets. **(A)** EpiTox-based identification of physiologically relevant off-target peptides for KVL and GVY. Sequence similarity thresholds were set to 4 (KVL, 9-mer) and 5 (GVY, 10-mer), yielding similar pools of 1,181 (KVL) and 1,012 (GVY) peptides. Each dot represents one off-target peptide, plotted by EpiTox Risk Score 1 (Bi-feature ranking score, x-axis) and EpiTox Risk Score 2 (Multi-feature ranking score, y-axis). Red dots indicate peptides in the Irreducible risk core (see Methods); grey dots indicate peptides not meeting either tier. The orange region marks the danger quadrant (both scores ¿ 0.5); 9.4% of GVY and 23.3% of KVL peptides fall within it. Numbers at the top of each panel indicate the count of grey and red peptides within the danger quadrant, respectively. See Methods for details on the scoring methodology and expression-tier definition. **(B)** Differential tumor presentation of MAGE-A4 KVL and GVY epitopes across PDX models. Presentation of HLA-A*02:01-restricted MAGE-A4-derived epitopes GVYDGREHTV and KVLEHVVRV were assessed by immunopeptidomics in patient-derived xenograft (PDX) models of non-small-cell lung cancer (NSCLC), head and neck squamous cell carcinoma (HNSCC), and breast cancer (BRCA). A red X indicates samples in which the respective peptide-HLA (pHLA) complex was not detected. **(C)** Structure-based prioritization of MAGE-A4 GVY over KVL for TCRm targeting. PDB structures 8FJA and 8FJB (23) of the GVY and KVL peptides, respectively, in complex with HLA-A*02:01. The HLA α1 helix is shown in red, the α2 helix in green, and the β2-microglobulin subunit in grey. Blue and red indicate regions of positive and negative electrostatic potential on the peptide surface, respectively. Maximum side-chain deviation of the peptide from the HLA anchor line is reported in Å.

#### Patient-derived xenograft immunopeptidomics reveals context-dependent stability of pHLA epitope presentation

Recently, it was reported that the median target density in solid tumor tissues is significantly higher for KVL than for GVY when assessed within the same tumor samples. Given that peptide presentation is inherently context dependent [25], we next evaluated epitope presentation across patient-derived xenograft (PDX) models representing diverse tumor entities using immunopeptidomics. In contrast to the higher abundance of KVL observed in samples in which both epitopes were co-detected, GVY was consistently identified across all analyzed tumors, whereas KVL was detected in only 50% of samples (Figure 2 B).

Collectively, these findings indicate a trade-off between target abundance and breadth of presentation. While KVL may offer higher epitope density in selected contexts, GVY demonstrates a more uniform and broader presentation profile across tumor models, which may translate into a higher fraction of patients eligible for targeting. This broader and more consistent presentation supports the translational relevance of GVY as a MAGE-A4-derived pHLA target.

#### Peptide topology and solvent exposure define TCRm-accessible pHLA binding surfaces

For TCRm antibody generation, the pHLA complex should present a peptide-dominant, highly exposed surface in which solvent-accessible peptide residues protrude from the HLA groove in a defined and stable conformation. This enables antibody engagement of multiple peptide-specific contact points rather than reliance on conserved HLA framework interactions, a prerequisite for true TCR-like specificity. Ideally, the peptide adopts a rigid, low-flexibility conformation, minimizing conformational heterogeneity that can compromise antibody binding.

Structurally, the peptide-derived surfaces were analyzed directly from the experimental cryo-EM structures of the two pHLA-Fab complexes (PDB 8FJA, MAGE-A4 GVYD-GREHTV; PDB 8FJB, KVLEHVVRV; [23]). GVYDGRE-HTV forms a broader and more distinct peptide-derived surface within the HLA groove, characterized by a pronounced central backbone bulge that generates an extended solvent-exposed interface. The maximum side-chain deviation from the HLA anchor line (defined by the P1 and C-terminal anchor Cα positions) is 13.1 Å for GVYDGREHTV, reaching its peak at the Arg6 guanidinium group (NH1), compared with 8.9 Å for KVLEHVVRV, whose maximum is reached earlier, at the Glu4 carboxylate (OE1). This difference, together with the broader profile of the GVYDGREHTV bulge, reflects a more extended and more centrally displayed epitope topology for the 10-mer peptide.

In addition, GVYDGREHTV displays multiple solvent-exposed, chemically diverse residues, including Asp4, Arg6, His8, and Thr9. Among these, Arg6 sits at the apex of the backbone bulge and represents a central interaction hotspot, while His8 and Thr9 provide additional C-terminal contact points. In contrast, KVLEHVVRV exhibits a shorter and chemically less differentiated peptide surface. Although residues such as Glu4, His5, and Arg8 are solvent-exposed, the interface is more hydrophobic in character, dominated by valine and leucine residues (five of nine positions), which provide less distinctive chemical information for antibody discrimination. Consequently, antibody binding to this complex is more likely to rely on conserved HLA surface features, which is suboptimal for selectivity and safety. Taken together, GVYDGREHTV presents a more pronounced peptide-dominant surface architecture, supporting improved suitability for highly selective TCRm antibody recognition.

In conclusion, integrating a favorable predicted off-target landscape, a structurally optimized epitope surface, and broad tumor-associated presentation across PDX models, we identified GVYDGREHTV as the preferred target peptide for TCRm development. These converging lines of evidence indicate higher clinical relevance and an increased probability of successful development of TCR-like TCRm antibodies, supporting prioritization of this epitope for down-stream discovery efforts.

### Systematic Identification of TCR-Mimic Antibodies Targeting MAGE-A4 pHLA Complexes

A critical concern in the development of TCR-mimic antibodies (TCRms) is the potential for cross-reactivity with off-target pHLA complexes. To systematically minimize this risk, we implemented a multi-tiered selection and screening strategy designed to identify maximally de-risked TCRm candidates, built on three sequential pillars (Figure 3 A). First, generated antibodies are subjected to HighSCORE microarray-based XScanning to assess peptide footprint geometry. Candidates are prioritized based on maximal CDR contacts preferentially engaging centrally located peptide residues emerging from the HLA binding groove, mimicking the interaction geometry of native TCRs with pHLA complexes, over contacts directed towards N- or C-terminal peptide residues or the HLA framework. Peptide-centric binding modes identified by this approach are subsequently validated by T2-cell-based alanine scanning, confirming the physiological relevance of the observed interaction in a cellular context. Second, antibodies passing this entry-level selection are screened against a curated panel of physiologically relevant, sequence-similar pHLA complexes predicted by EpiTox, using HighSCORE-based microarrays to systematically define target-unrelated interactions for each TCRm candidate. Third, TCRms exhibiting the lowest number of critical off-target interactions are advanced to EpiPredict analysis, a TCRm-centric machine learning framework trained on kinetic binding data derived from XScanning and candidate-specific off-target datasets [18]. EpiPredict predicts sequence-unrelated off-target pHLA complexes, and candidate antibodies are subsequently screened against this computationally defined panel to provide a comprehensive, proteome-wide assessment of potential cross-reactivity. If this three-tiered strategy yields more candidates than can be feasibly advanced, an additional filtering step assessing sequence liabilities, including potential post-translational modifications, can be applied to further narrow the candidate pool. Following this approach, we identified two TCRm candidates, VR-4 and VR-6, with superior off-target profiles, described in detail in the following section together with a comparison to a patent-derived optimized MAGE-A4-directed TCRm VR-58.

**Figure 3.**
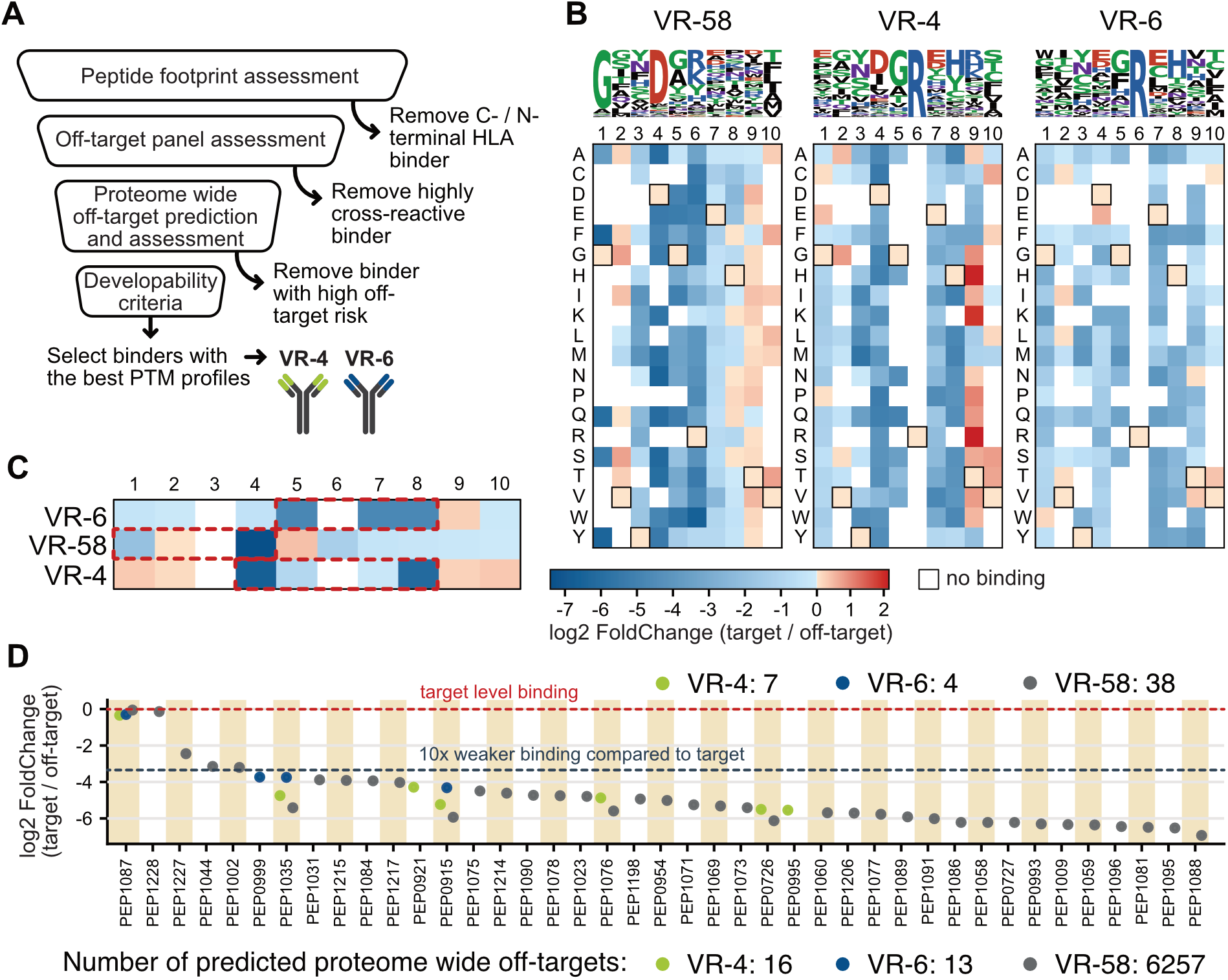
Systematic identification of TCR-mimic antibodies with superior off-target profiles. **(A)** Schematic workflow for the systematic identification of TCRm antibodies with superior off-target profiles. Target-specific antibodies from a TCRm library were sequentially screened by XScanning and T2-cell alanine scanning (peptide footprint assessment), EpiTox-predicted sequence-similar off-targets, and EpiPredict-based proteome-wide prediction of sequence-dissimilar off-targets. An optional additional filter assessing sequence liabilities, including potential post-translational modifications, can be applied if further narrowing of the candidate pool is required. Applying this workflow to MAGE-A4 identified VR-4 and VR-6 as lead TCRms with superior off-target profiles and no critical sequence liabilities. **(B)** XScanning logo plots and binding heatmap. Amino acid positions of the GVYDGREHTV decapeptide are indicated on the x-axis, single amino acid substitutions on the y-axis. Heatmap color intensity reflects relative binding, transformed as log2 fold change compared to the MAGE-A4 wild-type peptide (wild-type amino acids at each position are boxed): pink indicates binding similar to wild-type, colder blue colors indicate weaker binding, and warmer colors indicate stronger binding. White indicates no detectable binding. In the logo plots, letter size reflects the relative contribution of each amino acid position to TCRm binding. **(C)** Positional alanine scanning mutagenesis for epitope mapping. EC_50_ binding values for VR-4, VR-6, and VR-58 were measured against alanine-substituted MAGE-A4 peptide variants and log2-transformed relative to the wild-type peptide, using the same convention as in **(B)**. To exclude false negatives arising from insufficient peptide presentation, peptide loading was verified by β2-microglobulin staining post-loading (Figure S7). Numbers indicate positions in the MAGE-A4 peptide sequence that are replaced by alanine in this assay. Color coding as in **(B)**. Dashed red boxes indicate regions important for binding. **(D)** Off-target landscape overlap analysis. All off-target peptides identified by HighSCORE microarray screening for VR-4 (7 off-targets), VR-6 (4 off-targets), and VR-58 (38 off-targets) are plotted as log2-transformed affinity ratio relative to the MAGE-A4 wild-type peptide KD. VR-4 (green), VR-6 (blue), and VR-58 (orange) off-target peptides are displayed by peptide ID on the x-axis. An affinity ratio of less than 10-fold relative to the target, defined as the threshold for critical off-target risk, is indicated by a dashed line. Numbers of sequence-dissimilar off-target peptides predicted by EpiPredict for VR-4, VR-6, and VR-58 are shown beneath.

#### Peptide footprint mapping

As a first selection criterion, the peptide interaction profile of each TCRm candidate was determined using HighSCORE-based XScanning (Figure 3 B). Full binding kinetics were measured against a positional pHLA saturation library in which each position of the GVY decapeptide was individually substituted with all 19 alternative amino acids, generating 190 positional variants [18]. Positions at which substitutions retain wild-type-level relative binding are considered tolerated and thus of lesser relevance for TCRm contact. Conversely, positions at which substitutions result in reduced relative binding indicate dominant interaction relevance of the wild-type amino acid residue, reflecting potentially critical TCRm contact points. The three TCRm antibodies exhibit distinct binding interaction profiles with the MAGE-A4 pHLA complex. VR-58 displays a strong preference for N-terminally located glycine at position 1 and aspartate at position 4, indicating an N-terminally shifted epitope. In contrast, VR-4 and VR-6 critically require the centrally located arginine at position 6 for efficient binding. Both antibodies additionally show preferences for residues flanking Arg6: VR-4 favors aspartate at position 4, glycine at position 5, and histidine at position 8, while VR-6 similarly prefers wild-type glycine at position 5 and histidine at position 8, with threonine tolerated as a position-8 substitution with near wild-type affinity. Collectively, VR-4 and VR-6 fulfil the criterion for TCR-like peptide engagement, addressing centrally located residues with a broader peptide footprint, whereas the VR-58 epitope is shifted towards the N-terminus of the MAGE-A4 decapeptide. To confirm the relevance of the contact positions identified by microarray-based XScanning in a cellular context, T2 cells were pulsed with alanine-substituted variants of the MAGE-A4 decapeptide and EC50 binding of the three TCRm antibodies was determined (Figure 3 C). The alanine scan data was largely consistent with the XScanning results. For VR-58, a strong dependency on aspartate at position 4 was observed, with additional contributions from glycine at position 1 and arginine at position 6, as evidenced by increased EC50 values upon alanine substitution at the respective positions. For VR-4 and VR-6, the central importance of arginine at position 6 was confirmed, along with the contributing residues previously identified in the XScanning screen. Notably, loss of binding upon tyrosine-to-alanine substitution at position 3, observed for all binders, likely reflects a structural change in epitope configuration in a cellular context that is not captured by HighSCORE (see Structural analysis of VR-4, VR-6, and VR-58, below, and Figure S7, showing the loading data for T2 cells).

#### Off-target landscaping

To assess cross-reactivity, the three antibodies were screened in monovalent Fab format against 339 EpiTox-predicted peptides for the GVY epitope at a similarity threshold of four using HighSCORE-based microarray screening (Figure 1A). Potential off-targets identified for each TCRm are shown in Figure 3 D, including the affinity window relative to the target peptide as quantified by log2 target/off-target KD ratios. VR-58 bound 38 of the 339 peptides, whereas VR-4 showed significant binding to only 7 sequence-similar off-targets, and VR-6 displayed the most favorable profile with binding to only 4 off-target peptides (see Table S1 and Figure S1 for an annotated list of off-targets including tissue distribution and risk features). Off-target overlap among the three antibodies was minimal. PEP-1087 and PEP-1035, both derived from MAGE family members with high sequence similarity to MAGE-A4, were bound with high affinity by all three antibodies. Outside the MAGE family, VR-58 and VR-4 shared binding to PEP-0726, whereas VR-6 did not share any off-targets with the other two antibodies (Figure 3 E). PEP-0726 was pursued further in the following sections as an illustrative example of systematic off-target de-risking, given its wide tissue expression, high bi-feature similarity rank combining HLA-A*02 binding affinity and amino acid physicochemical properties, and experimental evidence supporting its physiological existence (Figure S1). With the exception of one peptide with very restricted tissue expression and low experimental evidence (PEP-0999), VR-6 bound exclusively to other MAGE family members, which can be considered de-risked given their limited expression in healthy tissues [26]. These findings are consistent with the distinct XScanning profiles of the three TCRms and suggest that central peptide binding footprints contribute to the more favorable sequence-similar off-target profiles of VR-4 and VR-6.

To confirm these findings in a cell-based system, VR-4, VR-6, and VR-58 were analyzed in IgG format for binding to selected peptide-pulsed T2 cells (Figure 4). Binding to the relevant sequence-similar off-target peptides was confirmed at least qualitatively for the respective VR binders (Figure 4A), consistent with the HighSCORE-based results. All three TCRms bound the MAGE-A4 wild-type peptide PEP-0705 with single-digit nanomolar EC50 values, with VR-4 and VR-58 IgGs displaying approximately fivefold higher apparent affinity than VR-6. Binding to PEP-1087 (MAGE-A8-derived) and PEP-1035 (MAGE-C2-derived) was confirmed for all three antibodies; however, complete dose-response curves for PEP-1035 could not be achieved for VR-4 and VR-58. A similar discrepancy was observed for PEP-0726, where VR-4 did not yield a complete dose-response curve despite tight monovalent binding in High-SCORE assays, whereas VR-58 did. These observations suggest that bivalent cell-based binding in IgG format may occasionally be subject to different constraints than monovalent Fab-based binding on the microarray, potentially attributable to higher local pHLA complex densities on the microarray surface relative to peptide-pulsed T2 cells. Importantly, all selected off-target peptides identified for VR-58 by High-SCORE screening could be verified in the T2-cell binding system, demonstrating that HighSCORE-based screening data are predictive for cell-based in vitro settings. In summary, XScanning and EpiTox-guided HighSCORE screening, followed by cell-based validation effectively stratify TCRm candidates by their off-target liability profiles, providing a robust early-stage risk assessment framework for candidate selection with VR-6 emerging as a very favorable anti-MAGE.

**Figure 4.**
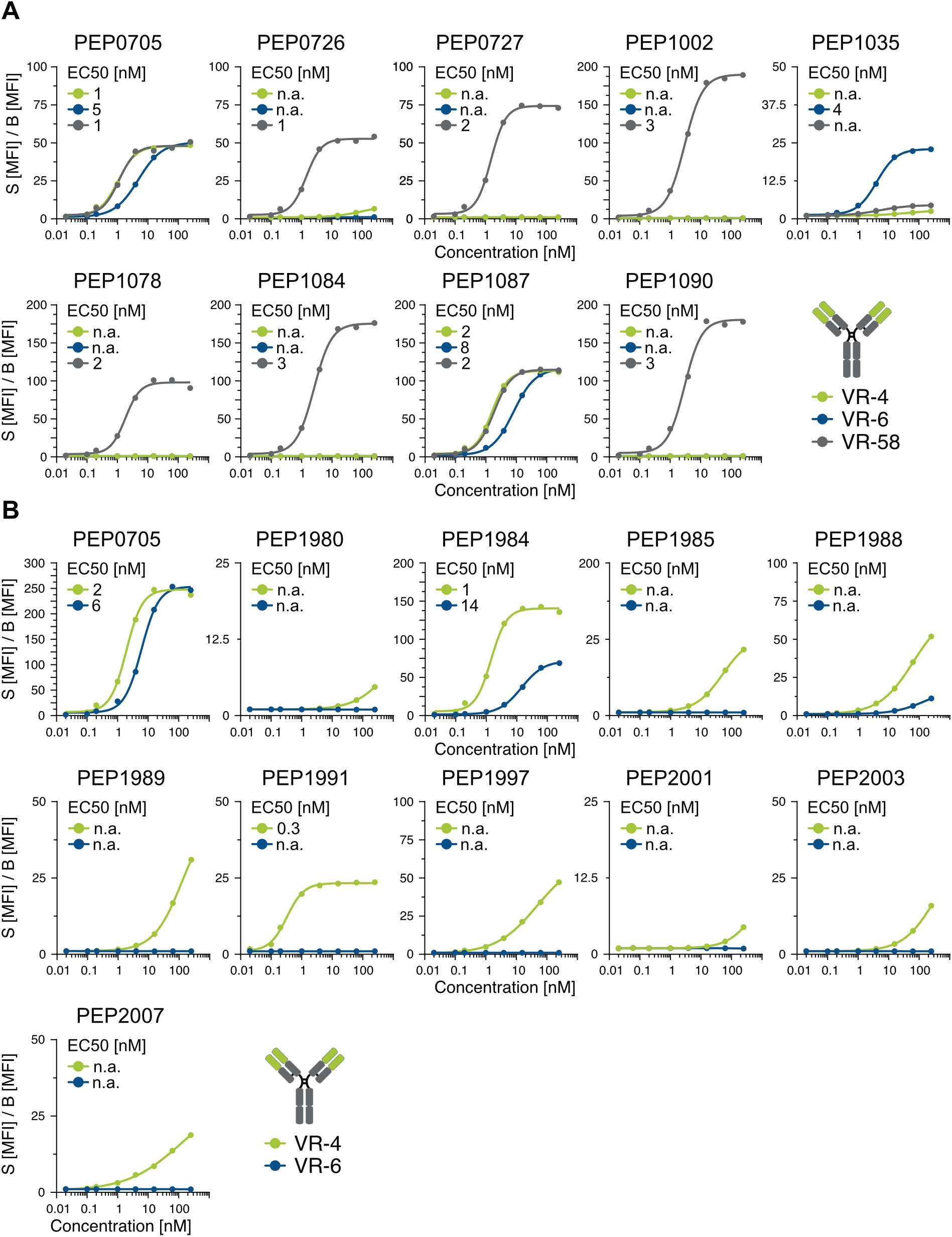
Binding of TCRm antibodies VR-4, VR-6, and VR-58 to MAGE-A4 target and selected off-target peptides in IgG format. **(A)** EpiTox-derived sequence-similar peptides. Binding dose-response curves of VR-4, VR-6, and VR-58 IgGs against T2 cells pulsed with 5 *µ*M of MAGE-A4 target peptide PEP-0705, MAGE family member peptides (PEP-1002, MAGE-B10-derived; PEP-1035, MAGE-C2-derived; PEP-1078, MAGE-B2-derived; PEP-1084, MAGE-A10-derived; PEP-1087, MAGE-A8-derived; PEP-1090, MAGE-B6-derived), and HighSCORE-identified off-target peptides PEP-0726 and PEP-0727. Antibody concentration (nM) is plotted on the X-axis (log10 scale) against signal-to-background (S(MFI)/B (Isotype control; MFI) on the Y-axis; the Y-axis maximum varies between peptides. EC50 values are indicated. VR-46, a non-targeting control derived from the anti-RSV antibody Motavizumab, served as negative control. **(B)** EpiPredict-derived sequence-dissimilar peptides. Binding dose-response curves of VR-4 and VR-6 IgGs against T2 cells pulsed with 100 *µ*M of the 10 experimentally confirmed EpiPredict-predicted binders. Antibody concentration (nM) is plotted on the X-axis (log10 scale) against signal-to-background (S(MFI)/B Isotype control(MFI)) on the Y-axis; the Y-axis maximum varies between peptides. EC50 values are indicated where dose-response curves could be completed. n.a., no binding signal or binding signal insufficient to complete dose-response titration.

#### Proteome wide off-target identification

While HighSCORE-based screening against sequence-similar peptides provides an early risk stratification frame-work, it does not capture potential binding to sequence-dissimilar peptides whose biophysical property features may yet complement the paratope of a given TCRm. To identify such sequence-dissimilar off-targets at proteome scale, kinetic data from XScanning and EpiTox-guided HighSCORE screening were used to train EpiPredict, our TCRm-centric predictive model for proteome-wide identification of high-affinity sequence-dissimilar binders [18]. As previously reported, VR-4 and VR-6 (therein referred to as antibody A and antibody B, respectively) were predicted by EpiPredict to bind 18 and 14 sequence-dissimilar peptides, respectively, of which 10 unique peptides, with some overlap between the two antibodies, could be confirmed by HighSCORE or T2-cell binding experiments. The annotated profiles of the confirmed sequence-dissimilar off-targets for VR-4 and VR-6 are shown in Figure S2. All predicted peptides were assessed for HLA-A*02 loading capacity (Figure S7) and for dose-responsive binding of VR-4 and VR-6 to HLA-A*02-loaded T2 cells. As shown in Figure 4 B, EC50 dose-response values could be achieved for only a small subset of predicted peptides (PEP-1984 and PEP-1991), and only at high loading concentrations of 100 *µ*M. The requirement for high peptide concentrations to achieve full dose response binding indicates weak HLA-A*02 binding affinity. Consequently, even where VR-4 or VR-6 displayed detectable EC50 binding to predicted sequence-dissimilar peptides, the underlying HLA-A*02 affinities are insufficient to support physiologically relevant pHLA surface presentation and thus preclude off-target cytotoxic activity under physiological conditions. All predicted sequence-dissimilar off-targets for VR-4 and VR-6 can therefore be considered de-risked.

Applying the same approach to VR-58, EpiPredict predicted 6,257 pHLA complexes as high-probability binders (Figure 3D; sequences available upon request). Experimental validation of this larger set was not pursued due to resource constraints; however, given the confirmation rate observed for VR-4 and VR-6, it is plausible that a substantial fraction of these predicted complexes represent genuine, clinically relevant off-targets that would require further characterization and de-risking, for example by T2-cell binding confirmation, further underscoring the unfavorable off-target risk profile of VR-58 already indicated by XScanning and EpiTox-guided HighSCORE screening of sequence-similar peptides. Also, none of the peptides predicted for VR-4 and VR-6 showed detectable binding by VR-58, even at high antibody concentrations (data available upon request), consistent with the distinct epitope recognition modes of VR-58 compared to VR-4 and VR-6 established by XScanning and alanine scanning (Figure 3 B, C). This cross-antibody specificity of EpiPredict predictions demonstrates that the model, when trained on TCRm-specific kinetic data from HighSCORE-based screening, captures the individual binding fingerprint of each TCRm with remarkable precision, discriminating binding interactions at the single amino acid level despite the inherently small target epitope of only 10 residues. This highlights the power of TCRm-specific kinetic training data in enabling EpiPredict to function as a TCRm-centric rather than a generic pHLA-binding predictive model.

Collectively, the ValidaTe workflow identifies VR-4 and VR-6 as TCRm candidates with peptide-centric, TCR-like binding behavior and favorable proteome-wide off-target profiles, supporting their advancement as lead candidates for clinical development. Notably, VR-6 emerged as the most favorable of the three candidates, displaying no detectable off-target risk outside the MAGE family.

### Structural analysis of VR-4, VR-6 and VR-58

To structurally validate the peptide-centric, TCR-like binding geometry inferred from screening and to establish the molecular basis for the favorable off-target profiles of VR-4 and VR-6 relative to VR-58, we determined cryo-EM structures of all three TCRm Fabs in complex with the MAGE-A4 GVYDGREHTV pHLA-A*02:01 complex at high resolutions of 2.2 to 2.4 Å (see supplementary Table S4), revealing the location of numerous ordered solvent molecules. Interaction energies are MOE-calculated estimates (MOE version 2024.0601). In every antibody-pHLA complex the peptide adopts a similar conformation, with a central backbone bulge displaying Arg6 as the apex residue (Figure 5 A). A conserved Tyr3(OH)-Arg6 hydrogen bond (2.7 Å in all three structures) stabilizes this bulge, consistent with the rigid, peptide-dominant epitope and the strong Tyr3 sensitivity observed in cellular assays. Superimposed on this shared scaffold, each antibody stabilizes a distinct set of secondary intra-peptide contacts: in VR-4 the Thr9 hydroxyl caps the C-terminus through a Thr9(OG1)-Val10 carboxylate hydrogen bond (2.7 Å) absent in the other two complexes; in VR-6 the His8 imidazole (ND1) hydrogen-bonds to the Arg6 main chain (2.9 Å), a contact only marginal in VR-4 and absent in VR-58; and in VR-58 the Asp4 side chain forms a short self-capping hydrogen bond to its own backbone amide (2.7 Å). Thus, each antibody selects and rigidifies a slightly different peptide micro-conformation around the common Tyr3/Arg6 core. VR-4 engages the peptide across positions 3-9 through an extensive, peptide-centered interface (20 antibody-peptide contacts; 6 hydrogen bonds including 3 buried ionic salt bridges). The dominant interaction is a CDR-H2 Glu51-Arg6 salt bridge (closest heavy-atom approach 1.7 Å), reinforced by a second heavy-chain acidic contact (Asp99-Arg6) and a Lys33-Asp4 salt bridge (Figure 5 B; VR-4). The calculated antibody-peptide energy is ≈-88 kcal/mol. Antibody contacts to the HLA α1/α2 helices are more limited (29 contacts; 9 hydrogen bonds, a single salt bridge; Figure 5 C; VR-4), the strongest being CDR-L2 Glu55-Arg66(α1) (2.8 Å); CDR-H3, -L1 and -L2 engage the α1 helix while CDR-H1/-H2 engage α2 (Figure 5 C; VR-4). The calculated antibody-HLA energy ≈-52 kcal/mol; total ≈-140 kcal/mol.VR-6 recognizes the same peptide surface (positions 3-9; 23 antibody-peptide contacts, 7 hydrogen bonds including one salt bridge), with the central CDR-H3 Glu99-Arg6 salt bridge (1.7 Å) as the dominant peptide contact (Figure 5 B; VR-6). The calculated antibody-peptide energy is ≈-52 kcal/mol. HLA engagement (35 helix contacts; 4 hydrogen bonds including one salt bridge and one arene contact) is led by a CDR-H2 Arg54-Glu167(α2) salt bridge (Figure 5 C; VR-6). The calculated antibody-HLA energy is ≈ −50 kcal/mol; total ≈ −102 kcal/mol. Uniquely among the three peptide structures, the His8 side chain forms a hydrogen bond to the central peptide Arg6 backbone chain. VR-58, in contrast, forms a smaller and more N-terminally restricted peptide footprint (core contacts at positions 4-6; 13 antibody-peptide contacts, 5 hydrogen bonds including one salt bridge) built around a CDR-L2 Asp50-Arg6 salt bridge (Figure 5 B; VR-58). The calculated antibody-peptide energy is ≈ −47 kcal/mol. Its HLA footprint is markedly larger and strongly placed toward the α2 helix, close to the A-pocket (41 helix contacts, α2:α1 6:1; 13 hydrogen bonds including one salt bridge), with CDR-L1/-L3 and CDR-H3 contacting α2 and the strongest helix contact formed by CDR-L1 Arg29-Glu155(α2) (Figure 5 C; VR-58). The calculated antibody-HLA energy is ≈ −49 kcal/mol; total ≈-96 kcal/mol. Here the Asp4 side chain of the peptide forms the shortest intramolecular hydrogen bond to its own backbone amide (2.7 Å), while the conserved Tyr3-Arg6 hydrogen bond is retained.

**Figure 5.**
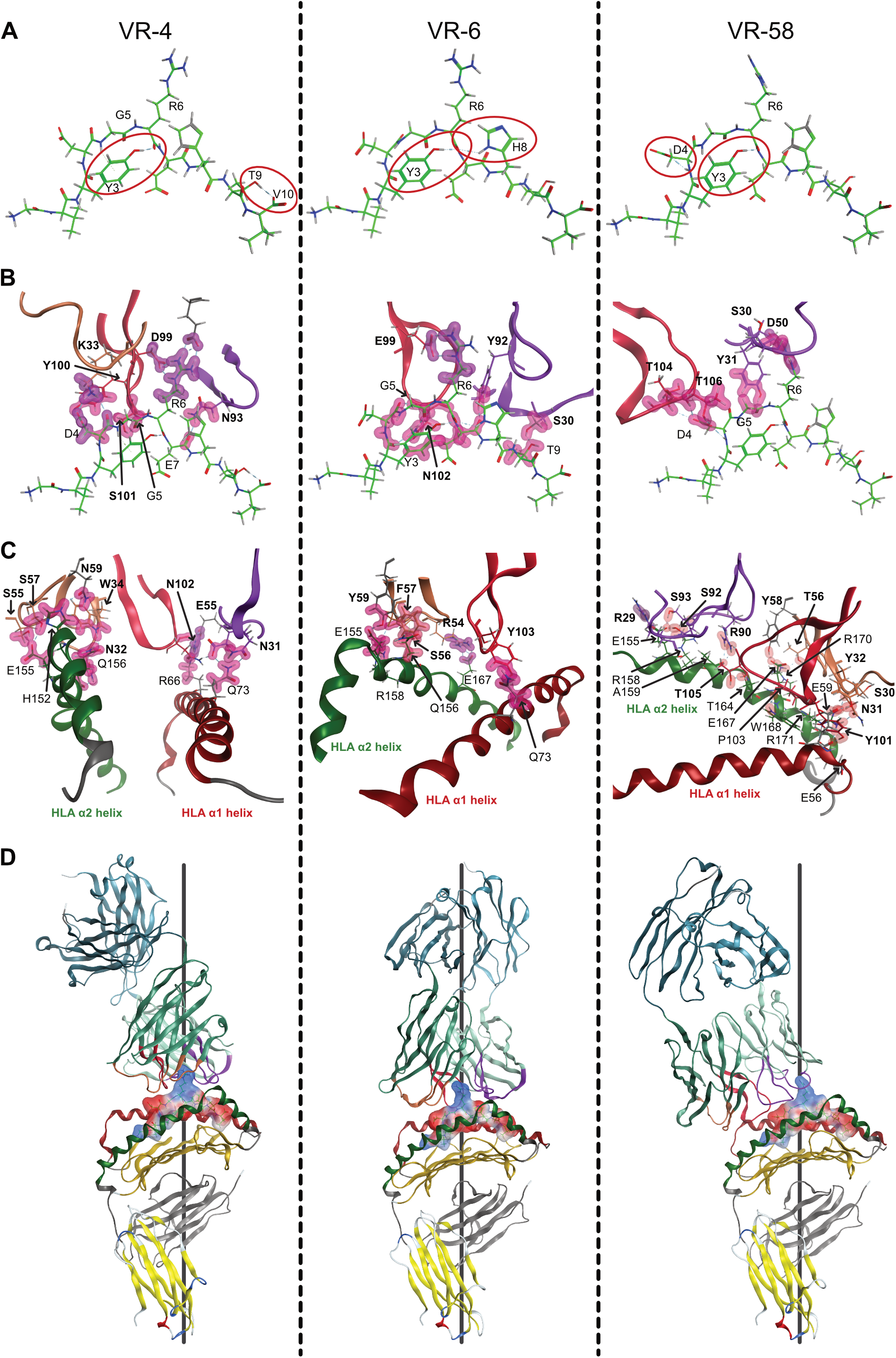
Cryo-EM structures reveal a shared peptide-recognition core with distinct binding geometries underlying the favorable off-target profiles of VR-4 and VR-6. **(A)** MAGE-A4 peptide (GVYDGREHTV) bound to VR-4, VR-6, and VR-58 (left to right; antibody and HLA omitted). Red circles mark intramolecular H-bonds: the conserved Tyr3-Arg6 bond (2.7 Å) at the Arg6 bulge in all three complexes, plus an antibody-specific secondary contact - Thr9-Val10 (VR-4), His8-Arg6 (VR-6), Asp4 sidechain to backbone amide (VR-58). Atom colors: carbon, green; nitrogen, blue; oxygen, red; hydrogen, grey (added computationally for visualization; not resolved in the cryo-EM density). **(B)** Antibody recognition of the peptide in the VR-4, VR-6, and VR-58 pHLA complexes (framework regions and HLA omitted). CDR-H1/H2/H3, brown/red ribbons; CDR-L1/L2/L3, purple; bold residues denote antibody, non-bold denote peptide/HLA (applies to **B, C**); H-bonds, red cloud; ionic contacts, purple cloud. VR-4 and VR-6 engage an extended peptide surface (positions 3-9) via CDR-H2/CDR-H3-Arg6 salt bridges; VR-58 shows a smaller, N-terminally restricted footprint (positions 4-6) via a CDR-L2-Arg6 salt bridge. **(C)** Antibody contacts to the HLA α1/α2 helices, with CDR-HLA contact residues shown as sticks; coloring as in **(B)**. VR-4 and VR-6 engage α1 and α2 in a balanced manner; VR-58 contacts concentrate on α2, proximal to the A/B-pocket. **(D)** Overall Fab-pHLA architecture: central, symmetric, TCR-like geometry of VR-4 and VR-6; relative to the A-pocket/α2-shifted mode of VR-58. Coloring as in **(B)**; HLA α1 helix, red; α2 helix, green; β2-microglobulin, grey. Blue/red indicate positive/negative electrostatic potential on the peptide surface. Antibody constant domains, though present in the cryo-EM map, showed lower local resolution and were not included in the deposited coordinates; they are retained in the figure for structural orientation only. Summary of structural features in table S3.

Comparison of the three interfaces reveals a common recognition principle and a clear geometric divergence (Figure 5 D). In all three antibodies the single strongest peptide contact is an acidic-residue salt bridge to the central Arg6, of similar calculated binding strength (≈ −32 to −37 kcal/mol), underscoring Arg6 as the dominant specificity determinant. However, VR-4 and VR-6 bind centrally and symmetrically over the peptide in a TCR-like manner, with balanced HLA α1/α2-helix engagement and most of their binding energy from peptide contacts, whereas VR-58 is shifted closer to the A-pocket toward the α2 helix, engages fewer peptide positions, and derives a larger share of its binding energy from conserved HLA surfaces, consistent with its less favorable off-target profile.

#### ParaPredict models of VR-4 and VR-6 accurately recapitulate the experimentally determined binding geometry

We previously developed the structure-prediction workflow underlying ParaPredict, an AI-assisted approach that generates in silico TCRm-pHLA complex models to guide antibody optimization, and reported models for VR-4 and VR-6 (referred to as antibody A and antibody B, respectively, in that publication) [18]. With experimental cryo-EM structures of both complexes now available, we assessed the accuracy of these models by superposing them onto the corresponding experimental structures. Both models faithfully reproduced the peptide-centric, TCR-like binding mode, including near-quantitative agreement for the dominant antibody-peptide and antibody-HLA salt bridges ((Figure S8; see supplementary text for complete structural comparison)). Differences were limited to specific features of the docking geometry, namely a single side-chain interaction in the VR-4 model and a modest global orientation shift in the VR-6 model, while the overall interaction network was preserved. Together, these findings demonstrate that ParaPredict generates structurally accurate models that faithfully capture the key molecular interactions observed in experimentally determined TCRm-pHLA complexes, supporting its use as a reliable starting point for interface analysis and off-target prediction. Because ParaPredict is designed as an iterative framework, experimental data generated throughout the ValidaTe workflow, including HighSCORE-based peptide XScanning, antibody CDR mutagenesis, phage display enrichment, and additional structural information, can be incorporated to progressively refine the predicted binding geometries and interaction interfaces, further improving the predictive performance of the platform.

### Functional Validation via TCRm-mediated cellular cytotoxicity

HighSCORE-based off-target screening of sequence-similar MAGE-A4-related peptides, combined with ML-based proteome-wide off-target prediction of sequence-dissimilar peptides, demonstrated a favorable off-target profile for VR-4 and VR-6 compared to VR-58. This was further supported by structural analysis, consistent with the larger and more centrally located interaction footprints of VR-4 and VR-6, which correlated with reduced off-target peptide binding. To substantiate the safety relevance of these findings, we next sought to verify a selection of physiologically relevant off-target peptides in a cellular context and to assess their capacity to mediate TCRm-dependent cytotoxicity.

To evaluate the ability of VR-4, VR-6, and VR-58 to mediate T-cell-redirected cytotoxicity, all three TCRm antibodies were reformatted into asymmetric 1+1 bispecific scFv-Fab-Fc T-cell engagers (TCEs). TCE purity was confirmed by SEC (Figure S3 A), and constructs were confirmed to retain binding to the MAGE-A4 target pHLA complex PEP-0705 and selected off-target peptides by flow cytometry using peptide-pulsed T2 cells (Figure 6 A); CD3 engagement via the anti-CD3 moiety was verified using primary T-cells, with all constructs exhibiting similar EC_50_ binding values (Figure S3 B).

**Figure 6.**
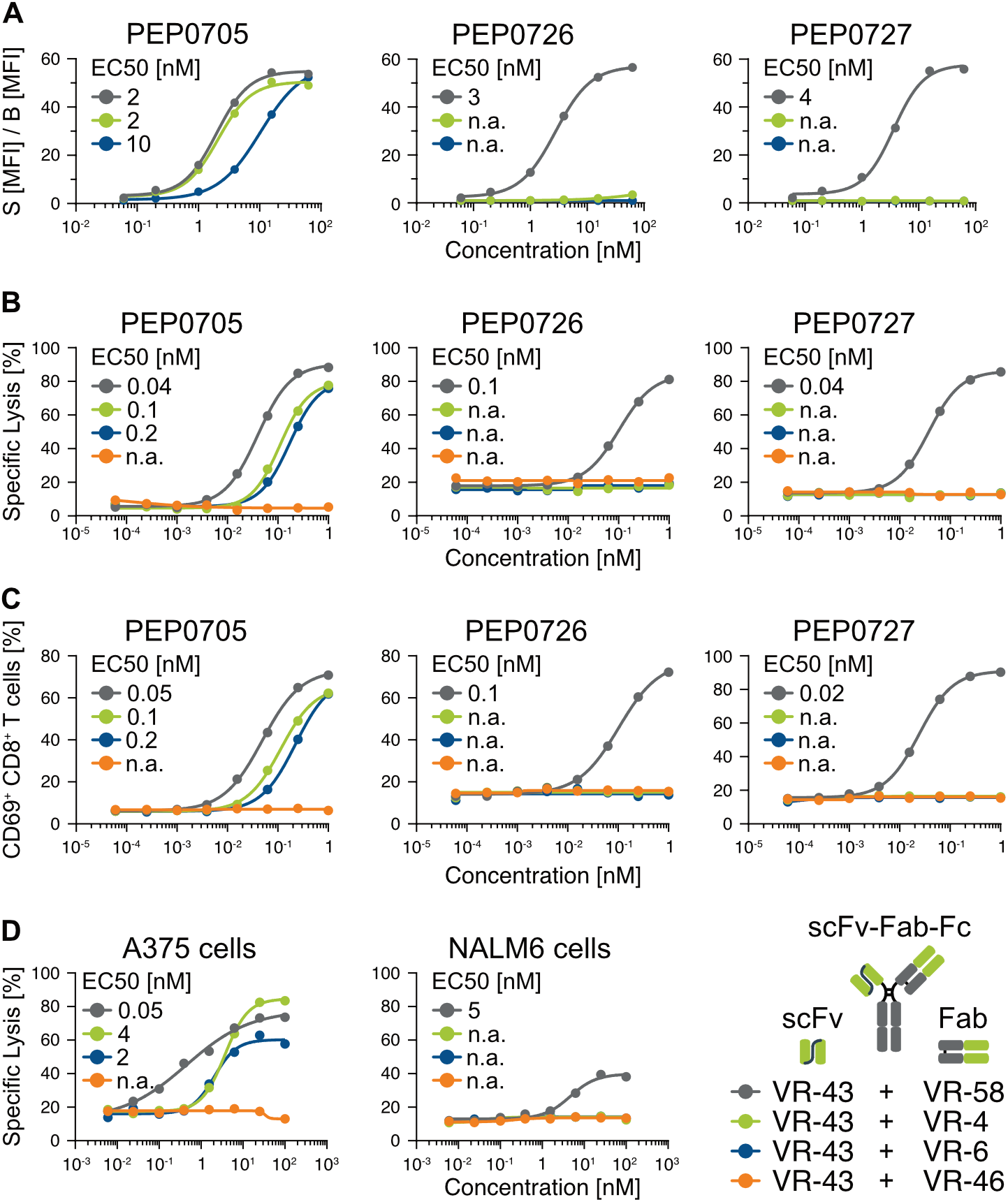
Functional validation of ValidaTe-identified off-target risk via TCRm-based T-cell engager cytotoxicity assays. Schematic representation of the bivalent scFv-Fab-Fc 1+1 format, indicating the anti-CD3-binding domain VR-43 on the scFv arm and the variable binding moieties on the Fab arm. Dot colors in the schematic correspond to the colors used in the dose-response curves. EC_50_ binding values for the indicated VR combinations were determined by flow cytometry using T2 cells pulsed with 5 *µ*M of the MAGE-A4 target peptide PEP-0705 or the off-target peptides PEP-0726 and PEP-0727. Antibody concentration (nM) is plotted on the x-axis (log10 scale) against signal-to-background (**A**; S(MFI)/B Isotype control(MFI)), specific lysis (% dead cells; **B** and **D**), or CD69+ cells (%; **C**) on the y-axis; EC_50_ values are shown. n.a., no binding signal. **(A)** EC_50_ binding values of VR-4-, VR-6-, and VR-58-based TCEs to primary T-cells and to T2 cells pulsed with 5 *µ*M of the MAGE-A4 target peptide PEP-0705 or the off-target peptides PEP-0726 and PEP-0727. VR-43 represents the isolated anti-CD3-binding moiety; VR-46 serves as a non-targeting control derived from the anti-RSV antibody motavizumab. **(B)** Specific lysis and **(C)** T-cell activation dose-response of VR-4-, VR-6-, and VR-58-based TCEs against T2 cells (4h) pulsed with 5 *µ*M of the MAGE-A4 target peptide PEP-0705 or the off-target peptides PEP-0726 and PEP-0727. **(D)** TCE-mediated killing of HLA-A02^+^/MAGE-A4^+^ A375 melanoma (24 h) and HLA-A02^+^/MAGE-A4^−^ NALM-6 leukemia cells (4 h).

TCE-mediated killing of T2 cells pulsed with target or selected off-target peptides was subsequently assessed (Figure 6 B). Consistent with the binding data, all three TCEs induced potent killing of T2 cells loaded with the MAGE-A4 target peptide, accompanied by robust T-cell activation as evidenced by CD69 upregulation after 4 hours (Figure 6 C), and IFN-γ secretion after 24h (Figure S3C). Notably, despite VR-4 displaying approximately fivefold higher binding affinity for MAGE-A4-loaded T2 cells relative to VR-6, both TCEs mediated similar cytotoxic efficacy, indicating that binding affinity alone does not determine lysis potency and that additional context-specific features contribute to efficient target engagement to mediate cytolysis. In contrast, when T2 cells were pulsed with selected off-target peptides PEP-0726 and PEP-0727 at 5 *µ*M, only VR-58-based TCEs induced off-target cell killing and associated T-cell activation. Notably, PEP-0726, identified as a shared off-target binder of both VR-4 and VR-58 using HighSCORE, induced potent cytotoxicity only in the context of VR-58, demonstrating that monovalent binding affinity alone does not necessarily predict functional risk in a cellular context. These findings confirm that off-target peptides identified by the ValidaTe screening cascade can mediate T-cell-dependent cytotoxicity and therefore represent a potential risk for clinical development.

To assess off-target risk under more physiologically relevant conditions, TCE-mediated killing was evaluated using HLA-A*02-positive A375 melanoma cells, which present the MAGE-A4 GVY epitope as a naturally processed pHLA complex [27], and MAGE-A4-negative NALM-6 leukemia cells. In contrast to the supraphysiological peptide densities generated by 5 *µ*M peptide pulsing of T2 cells, this experimental setting reflects endogenous antigen presentation at physiological pHLA surface densities. Endogenous target pHLA presentation in A375 cells and its absence in NALM-6 cells were additionally confirmed and Western blotting using MAGE-A4-specific antibodies (Figure S5 A) and by immunopeptidomics (Figure S5 B-E). As shown in Figure 6 D, all three TCEs efficiently mediated killing of A375 cells, accompanied by IFN-γ release after 24 h and CD69 upregulation in CD8-positive T-cells after 4 h (Figure S4 A), with VR-58 exhibiting the greatest potency as indicated by a lower EC50. In contrast, only VR-58-based TCEs induced killing of target-negative NALM-6 cells (Figure 6 D), again accompanied by IFN-γ release (Figure S4 E).

Taken together, these results demonstrate the superior on-target specificity of VR-4 and VR-6 compared with VR-58 under both non-physiological peptide-pulsing and physiologically relevant endogenous antigen presentation conditions, supporting their advancement as lead TCRm-based T-cell engager candidates.

### Exploiting Distinct Off-target Fingerprints to Improve TCRm Selectivity

Using the ValidaTe platform, we identified MAGE-A4 GVY-specific TCRm antibodies with favorable off-target profiles. However, even a limited number of off-target interactions may constitute a safety liability by mediating unintended T-cell activation and cytotoxicity, underscoring the need for rigorous off-target assessment during preclinical development. Notably, subtle differences in pHLA recognition resulted in largely distinct off-target landscapes. This was exemplified by VR-6 and VR-58, which shared only peptides derived from homologous MAGE-A family members, representing anticipated and biologically de-risked cross-reactivities. These findings indicate that minor variations in target peptide engagement can profoundly influence off-target recognition despite similar target specificity. Based on these observations, we hypothesized that TCRms exhibiting non-overlapping off-target repertoires could be advantageously combined within a single therapeutic modality. Specifically, incorporation of two target-specific TCRm binding domains into a trivalent T-cell engager may increase avidity toward the target pHLA complex through cooperative binding, whereas most off-target pHLA complexes, recognized by only one of the two TCRms, would not benefit from such avidity effects. We term this design concept WiFi (Widened Fingerprint). The WiFi hypothesis proposes that combining TCRms with distinct off-target fingerprints while maintaining convergent target recognition may increase the probability of productive target engagement while reducing the contribution of individual off-target interactions. In the following sections, we experimentally evaluated whether this avidity-based design principle can improve the selectivity of TCRm-based T-cell engagers.

#### Combinatorial pairing of TCRm binders with divergent off-target profiles minimizes off-target reactivity while preserving target binding

To evaluate whether combining anti-MAGE-A4 TCRms with distinct off-target reactivity profiles could reduce overall off-target binding while preserving target specificity, we assessed the interaction of anti-MAGE-A4 TCRm pairs with selected off-target peptides using peptide-pulsed T2 cells. For this purpose, we introduced an additional anti-MAGE-A4 TCRm, VR-57, originally developed as a CAR-T binder with high specificity [23, 28]. Similar to VR-4 and VR-6, VR-57 displays high-affinity binding to MAGE-A4-pulsed T2 cells (Figure S6 A) and engages MAGE-A4 via more centrally located peptide residues within the pHLA complex (Figure S6 B). Consequently, VR-57 exhibits a distinct off-target profile that does not overlap with VR-58, except for PEP-1087 (MAGE-A8-derived), a feature it shares with VR-4, VR-6 and VR-58 (Figure S6 B). VR-4, VR-6, and VR-57 were each combined with VR-58 in the same bivalent scFv-Fab-Fc 1+1 format used in the cytotoxicity assays, with VR-58 held constant on the scFv arm and VR-4, VR-6, or VR-57 introduced on the Fab arm. All constructs were purified to at least 95% monomer content (Figure S3 A). Binding to the MAGE-A4 target peptide (PEP-0705) and to the off-target peptides PEP-0726 and PEP-0727 was assessed by flow cytometry using peptide-pulsed T2 cells. As shown in Figure 7, EC_50_ binding values for PEP-0705 remained largely unchanged upon replacing the VR-58 Fab arm with VR-4, VR-6, or VR-57, whereas marked differences emerged for the off-target peptides. Substituting the VR-58 Fab arm with VR-4 reduced off-target binding 3-to 15-fold, while substitution with VR-6 or VR-57, the two binders with the most favorable off-target profiles, resulted in near-complete abrogation of off-target peptide binding, with no measurable dose-response even at high antibody concentrations, resembling the binding behavior of combining VR-58 with the non-pHLA-targeting control VR-61 (derived from the anti-RSV antibody Palivizumab). Importantly, this reduction in off-target binding was achieved while high-affinity binding to the MAGE-A4 target peptide was maintained.

**Figure 7.**
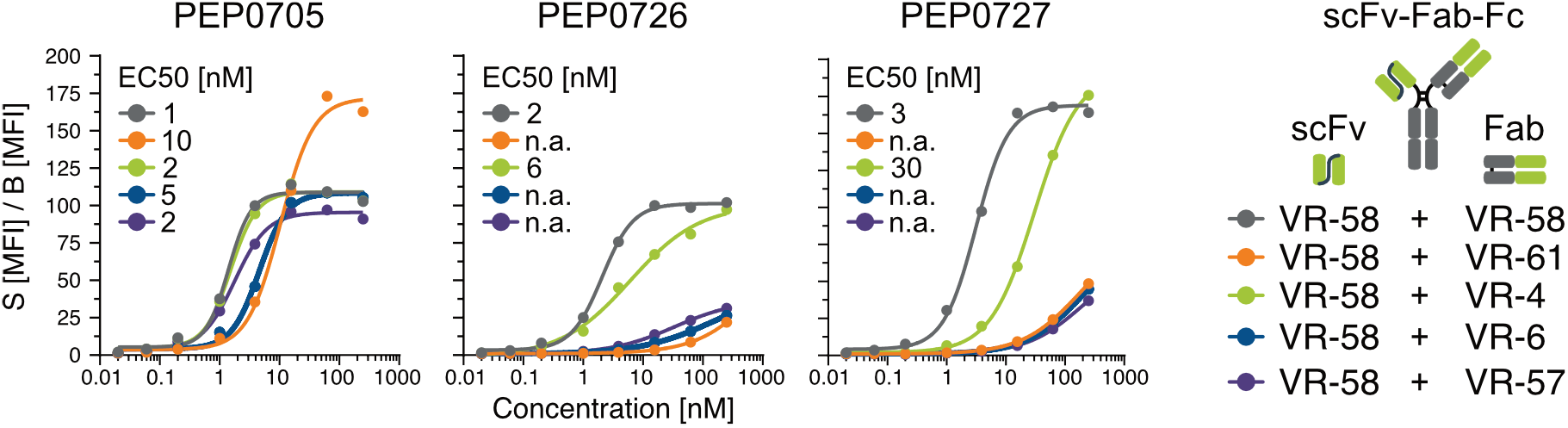
Combinatorial TCRm pairing reduces off-target peptide binding without compromising target affinity. Schematic representation of the bivalent scFv-Fab-Fc 1+1 format indicating VR-58 on the scFv arm and other variable binding moieties on the Fab arm. Dot colors in the schematic correspond to the colors used in the dose-response curves. EC_50_ binding values for the indicated VR combinations were determined by flow cytometry using T2 cells pulsed with 5 *µ*M of MAGE-A4 target peptide PEP-0705 or off-target peptides PEP-0726 and PEP-0727. Antibody concentration (nM) is plotted on the X-axis (log10 scale) against signal-to-background (S(MFI)/B(MFI)) on the Y-axis; EC_50_ values are shown. n.a., binding signal insufficient to complete dose-response titration.

#### Combinatorial pairing of TCRm binders with divergent off-target profiles minimizes off-target mediated cytotoxicity while preserving target-mediated

To evaluate the translational applicability of the WiFi concept, we next investigated whether the reduction in off-target binding achieved by combining TCRms with non-overlapping off-target profiles translates into reduced off-target-mediated cytotoxicity. To this end, constructs from the preceding binding experiments were re-engineered into trivalent 2+1 TCEs in a single-chain diabody-Fab-Fc (scDb-Fab-Fc) format incorporating an anti-CD3 binding domain for T-cell recruitment. In this format, VR-58 was retained on the Fab arm, as placing VR-58 on the scDb arm compromised construct stability. All constructs were purified to at least 95% monomer content to minimize the risk of non-specific T-cell activation (Figure S3 D); VR-4-containing constructs could not be purified above this threshold and were therefore excluded from further analysis. The remaining constructs were analyzed for binding to primary T-cells (Figure S3 E) and to T2 cells loaded with the MAGE-A4 target peptide PEP-0705 or the off-target peptides PEP-0726 and PEP-0727 (Figure 8; Binding).

**Figure 8.**
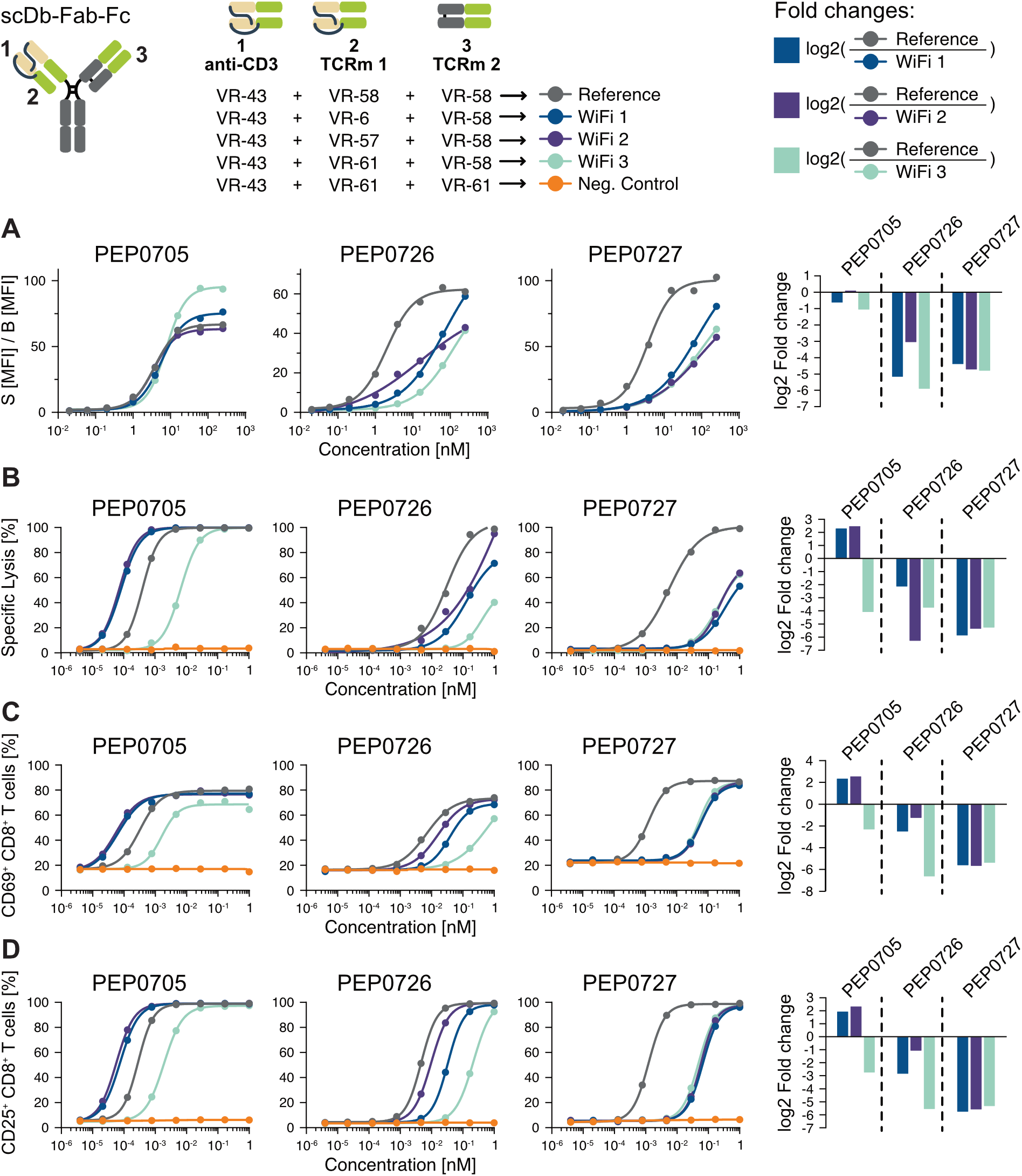
Combinatorial TCRm pairing strongly diminishes off-target-mediated T-cell redirected cytotoxicity while preserving target cell killing. Schematic representation of the trivalent scDb-Fab-Fc 2+1 format. The anti-CD3 binding moiety (brown) is positioned on the scDb arm (position 1), together with the anti-MAGE-A4 binding domains VR-6, VR-57, or VR-58, or the non-binding CTRL antibody VR-61 (derived from Palivizumab) (position 2, green). On the Fab arm (position 3), VR-58 or the VR-61 Palivizumab control is placed. The WiFi VR combinations (WiFi 1-3), in which VR-58 on the Fab arm is replaced by alternative anti-MAGE-A4 VRs, are color-indicated. VR-61 combinations on both the scDb and Fab arms served as the negative control (orange), whereas VR-58 combinations on both arms served as the reference (grey). **(A)** Binding, **(B)** T-cell-mediated cytotoxicity, and T-cell activation as measured by **(C)** CD69+ expression at 4 hours and **(D)** CD25+ expression at 24 hours were assessed using T2 cells pulsed with 5 *µ*M of the MAGE-A4 target peptide PEP-0705 or the off-target peptides PEP-0726 and PEP-0727, in the presence of donor-derived primary PBMCs. Results are expressed as log2 fold change relative to the VR-58/VR-58 reference. EC_50_ values for binding, killing, and T-cell activation were determined for each VR combination; where a full dose-response could not be achieved, values were interpolated. Negative log2 fold change values indicate diminished binding, killing, or activation relative to the reference.

All constructs bound primary T-cells and PEP-0705-pulsed T2 cells with similar EC50 values, with the exception of the VR-61/VR-61 control construct, which lacks an anti-MAGE-A4 binding domain. Off-target peptide binding to PEP-0726 and PEP-0727 was markedly reduced when VR-58 on the scDb arm was replaced by either the unrelated VR-61 domain or the MAGE-A4-targeting binders VR-6 or VR-57, consistent with the binding data obtained in the scFv-Fab-Fc 1+1 format.

Trivalent TCEs were subsequently evaluated in T-cell-mediated cytotoxicity assays using peptide-pulsed T2 cells and donor-derived primary PBMCs (Figure 8; specific lysis). After 24 hours, all constructs induced potent killing of PEP-0705-pulsed T2 cells, with the exception of the VR-61/VR-61 control. Notably, the VR-6/VR-58 and VR-57/VR-58 constructs were approximately fivefold more potent than the VR-58/VR-58 parental construct despite similar EC50 binding values, suggesting that factors beyond binding affinity contribute to cytotoxic potency. Consistent with this, T-cell activation, as measured by CD69 and CD25 upregulation in CD8-positive cytotoxic T-cells, was more pronounced for the VR-6/VR-58 and VR-57/VR-58 constructs (Figure S8; T-cell activation), indicating a contribution of the binding interface’s structural features to T-cell engagement. In contrast, TCE-mediated cytotoxicity against off-target peptide-pulsed T2 cells was markedly reduced upon replacement of the scDb-arm VR-58 moiety with VR-6 or VR-57, with correspondingly elevated EC50 values for T-cell activation.

Taken together, these results demonstrate that combining TCRms with distinct off-target binding profiles within the WiFi framework reduces off-target-mediated cytotoxicity while preserving, and in some cases enhancing, on-target activity. The data suggest that pairing binders with complementary recognition properties dilutes individual off-target interactions without compromising productive engagement of the target pHLA complex. This avidity-based strategy provides a rational approach to improving the selectivity and safety of TCRm-based T-cell engagers and may ultimately expand their therapeutic window.

## DISCUSSION

### Predictive de-risking enables safer pHLA-targeted therapeutics

Our findings address a major challenge in pHLA-targeted therapeutics: the need for predictive off-target screening technologies capable of de-risking soluble TCR- and TCRm-based T-cell engagers, as well as cell therapies such as TCR-T and CAR-T, before first-in-human studies. This need is particularly acute because conventional animal models poorly predict human pHLA-directed toxicity [29], while current de-risking strategies are either target-centric, largely ignoring the properties of the targeting molecule [15, 30], or too resource-intensive to support early-stage candidate discovery [27, 31]. ValidaTe addresses these limitations through an integrated workflow that enables rational decision-making across five complementary stages of pHLA therapeutic development: (i) selection of pHLA targets with favorable presentation, peptide topology, and predicted off-target landscapes; (ii) identification of peptide-dominant TCRm binders with TCR-like recognition properties; (iii) proteome-wide prediction and functional validation of off-target liabilities; (iv) therapeutic engineering strategies that further mitigate residual off-target risk while preserving on-target potency; and (v) orthogonal translational safety assessment to resolve residual uncertainties prior to clinical translation. These stages are designed to progressively reduce uncertainty, with increasingly resource-intensive approaches reserved for the progressively smaller number of candidates that advance through the workflow. Using MAGE-A4 as a proof-of-concept, we demonstrate how this framework supports the discovery of TCRm-based therapeutics with improved safety profiles while maintaining potent redirected cellular cytotoxicity. Extending this concept, the WiFi (Widened Fingerprint) principle illustrates how complementary recognition fingerprints can further expand the therapeutic window through rational therapeutic engineering.

### Level 1 - Target selection: Rational pHLA target selection requires integration of presentation, off-target space, and peptide topology

An optimal pHLA target for TCRm antibody development should combine broad tumor presentation, a limited predicted off-target space, and a peptide topology favoring peptide-dominant recognition. In this context, GVYDGRE-HTV emerged as a more favorable MAGE-A4 target than KVLEHVVRV. Although KVL reached higher abundance in selected tumor samples [22], GVY was detected more consistently across PDX models, suggesting broader patient coverage. Moreover, GVY exhibited a substantially smaller predicted high-risk off-target space, whereas KVL was associated with numerous sequence- and biophysically similar candidate off-targets, including many peptides expressed in healthy tissues (Figure 2 A). Supporting the predictive value of this metric, screening of EpiTox-ranked off-target peptides recently uncovered more than 20 previously unrecognized cross-reactive peptides for an anti-MAGE-A3 TCRm [20], consistent with our identification of more than 30 novel off-target binders for VR-58. Together, these findings suggest that the extent of the predicted high-risk off-target space may serve as a useful surrogate for cross-reactivity risk.

Structural analyses further supported the prioritization of GVY. Compared with KVL, GVY presents a more protruding and chemically diverse peptide surface with multiple solvent-exposed contact residues, favoring peptide-focused recognition. In contrast, the flatter, more hydrophobic surface of KVL may bias antibody binding toward conserved HLA features rather than the peptide itself, potentially reducing selectivity, consistent with structure-based analyses of pHLA recognition and TCR engagement [32]. This consideration may be less relevant for native TCRs, which interrogate a broader structural context of the pHLA complex [2, 33]. Indeed, KVL-targeting TCR-based T-cell engagers have recently demonstrated potent antitumor activity with a favorable off-target profile [22]. Collectively, these findings indicate that peptide topology should be considered alongside tumor presentation and predicted off-target space when prioritizing pHLA targets for TCRm discovery. Within this framework, GVYDGREHTV emerges as the preferred MAGE-A4 target, although comprehensive experimental validation remains essential to fully establish its safety profile.

### Level 2 - Binder selection: Peptide-dominant recognition features define selective TCRm binding

TCRm antibodies should preserve key features of native TCR recognition of pHLA complexes, including diagonal docking, a peptide-centered footprint, and balanced engagement of peptide and HLA surfaces [2, 34]. However, diagonal docking alone is insufficient to define TCR-like recognition properties; rather, specificity depends on the overall contact topology and the relative contribution of peptide versus HLA contacts [35, 36]. Excessive reliance on conserved HLA surfaces or incomplete peptide engagement may reduce epitope discrimination and increase cross-reactivity [37]. Together, these observations suggest that low-cross-reactivity TCRms require peptide-dominant recognition with limited dependence on conserved HLA features.

Guided by these principles, our selection strategy, combining HighSCORE-based XScanning with T2 cell binding confirmation, identified two TCRm antibodies, VR-4 and VR-6, that recapitulate peptide-dominant recognition features of native TCRs. Both antibodies engage an extended peptide footprint (positions 3-9), positioned over the central peptide arginine (Arg6; CDR-H2, H3 in VR-4, CDR-H3 in VR-6), together with balanced engagement of both HLA α1 and α2 helices. VR-58, despite similar overall antibody-peptide and antibody-HLA binding energies, engages a markedly narrower peptide footprint (positions 4-6 only) and binds the HLA helices asymmetrically, contacting α2 near the A-pocket roughly six times more frequently than α1. This narrow, one-sided docking geometry, rather than a difference in binding energy, underlies the less favorable off-target profile of VR-58 relative to VR-4 and VR-6.

Our findings are consistent with the framework proposed by Raybould et al. [36], suggesting that TCR-like recognition is governed by contact topology and interaction distribution rather than docking geometry alone. They also align with the work of Householder et al. [38], who used deep-learning-guided de novo design to generate rigid TCR-mimic scaffolds that preferentially engage exposed peptide residues while reducing off-target binding. Notably, the substantial sequence diversity between VR-4 and VR-6 [18] indicates that our selection strategy enriches for structural and functional recognition principles rather than specific CDR sequence motifs.

Despite there is no direct antibody contact to peptide position 1 in any of the three structures, the XScanning data show a markedly stronger preference for Gly at this position for VR-58 than for VR-4 or VR-6. This is consistent with the narrow peptide footprint of VR-58 and its closer paratope proximity to this region, which leave it with less steric and energetic tolerance for a bulkier P1 side chain.

Although the single strongest salt bridge to Arg6 shows a similar calculated binding energy across all three antibodies (≈ −32 to −37 kcal/mol), the surrounding recognition network differs fundamentally in chemical character. In VR-4 and VR-6, the guanidinium group is engaged by an extensive, side chain-specific network, including bidentate salt bridges to both terminal nitrogens and, in VR-4, an additional Trp34 cation-*π* cage, that requires the specific planar, delocalized charge of the arginine side chain and cannot be satisfied by other residues. In VR-58, by contrast, only a single acidic side chain (Asp50) forms a salt bridge, while most surrounding contacts are backbone carbonyl/amide groups of the CDR-L1 loop, which are chemically more promiscuous acceptors/donors. This more backbone-mediated, less side chain-specific recognition mode plausibly explains the greater tolerance for non-Arg substitutions at position 6 observed for VR-58 in the XScanning data.

Structural analyses further revealed a shared molecular basis for peptide recognition across the identified antibodies. An intact Tyr3 is required because its hydroxyl group forms a direct hydrogen bond to the Arg6 backbone carbonyl (conserved in all three structures), holding the central Arg6 residue in the peptide’s characteristic bulged conformation presented by HLA-A*02:01. This structural constraint explains the strong sensitivity to Tyr3 substitutions observed in cellular assays, whereas substitutions are better tolerated in HighSCORE-based measurements, likely reflecting the increased stability of the engineered HLA-A*02:01 variant used in this assay [39].

Collectively, these findings demonstrate that functional peptide-centric screening can preferentially identify TCRms that recapitulate peptide-dominant recognition features of native TCRs despite substantial sequence diversity. This supports a general strategy for discovering highly peptide-focused pHLA binders with improved specificity and reduced off-target liability.

### Level 3 - Integrated off-target de-risking: Proteome-wide prediction and functional validation of off-target liabilities

Our three-tiered discovery strategy identified VR-4 and VR-6 as highly specific anti-MAGE-A4 TCRms with TCR-like, peptide-centric recognition. Following sequential de-risking by EpiTox-HighSCORE screening and EpiPredict proteome-wide off-target prediction, only a small number of biologically plausible candidate off-targets remained, with VR-6 exhibiting the cleanest predicted profile. These predictions translated into functional outcomes: whereas VR-58 mediated potent cytotoxicity against both off-target peptide-loaded T2 cells and MAGE-A4-negative primary tumor cells, VR-4 and VR-6 showed no detectable off-target killing. Notably, VR-58 also displayed considerably greater cytotoxicity than VR-4 or VR-6 against A375 cells, potentially reflecting additional off-target interactions within this cellular context. Together, these findings demonstrate that an integrated de-risking strategy combining computational off-target prediction, high-throughput pHLA screening, and functional cellular validation can identify liabilities that translate into biologically relevant off-target cytotoxicity. This integrated approach highlights the importance of assessing both molecular recognition properties and functional consequences early during TCRm development to maximize the therapeutic window of pHLA-targeted therapeutics.

### Level 4 - Therapeutic engineering: The WiFi principle: engineering complementary off-target recognition fingerprints to expand therapeutic windows

These observations also inspired a complementary strategy to mitigate residual off-target toxicity at the therapeutic format level. Scheinberg and colleagues recently demonstrated that trivalent TCRm-based T-cell engagers targeting two unrelated tumor-specific pHLA complexes can reduce antigen escape while maintaining potent antitumor activity [40]. Building on this concept, we instead combined TCRms with complementary binding modes and non-overlapping off-target fingerprints. We hypothesized that combining TCRms with complementary recognition fingerprints would preserve avidity-driven target engagement while reducing effective affinity toward individual off-target peptides, which are recognized by only one component of the therapeutic. Consistent with this model, combining VR-58 with either VR-6 or VR-57 largely abolished VR-58-mediated off-target cytotoxicity while maintaining robust target-specific redirected killing. These findings establish the WiFi (Widened Fingerprint) principle as a new design strategy for TCRm-based therapeutics, demonstrating that off-target risk can be mitigated not only through antibody selection but also through rational therapeutic engineering. More broadly, this concept may provide a general framework for increasing the therapeutic window of pHLA-targeted biologics without compromising antitumor efficacy.

### Toward Level 5 - Translational safety assessment: Orthogonal validation of residual off-target liabilities

The molecular basis of the observed off-target toxicity of VR-58 remains to be elucidated. Although the gene encoding the candidate off-target peptide PEP-0726 (THBS3) is expressed in A375 cells (¿10 nTPM) [41], transcript abundance alone cannot establish peptide presentation and requires independent validation at the pHLA level. In a recent study, Marrer-Berger et al. demonstrated that mass spectrometry-based profiling of the TCRm-specific interactome provides a powerful downstream approach to identify HLA-presented off-target peptides in relevant cellular contexts [27]. A375 cells, which have been used as a rapid proxy system for establishing nonclinical PK/PD relationships for a MAGE-A4-targeting TCRm-based TCE due to similar EC_50_ potency to tumor organoid models [31], may therefore provide a relevant platform for such analyses. Although interactome profiling is not suitable for early antibody discovery, it represents a complementary final layer of the ValidaTe workflow. Following computational prediction, high-throughput screening, functional cytotoxicity assessment and therapeutic engineering, orthogonal proteomic characterization together with ex vivo primary human tissue testing can resolve residual uncertainties in peptide presentation and off-target biology before clinical translation.

Together, our study establishes ValidaTe as a platform for the rational development of pHLA-targeted therapeutics. Rather than treating off-target assessment as a late-stage safety exercise, ValidaTe integrates target prioritization, binder discovery, proteome-wide off-target prediction, therapeutic engineering, and orthogonal translational safety assessment into a unified development framework. By combining AI-based prediction, structural modeling, high-throughput pHLA screening, and functional validation, this iterative workflow enables systematic identification and mitigation of off-target liabilities while preserving therapeutic potency. Orthogonal proteomic characterization may further complement this framework as a final layer of safety evaluation prior to clinical translation.

## MATERIAL AND METHODS

### Molecular design, cloning, and production of TCRm-based mono-, bi-, and trivalent antibodies

Variable region sequences of VR-4 and VR-6 (derivable from the deposited PDB coordinates of the respective cryo-EM structures; see Data availability) and VR-58 (derived from WO2023110918A1; SEQ ID NO:54 [VH] and SEQ ID NO:55 [VL]), VR-57 [28], as well as the control antibodies VR-46 (motavizumab) and VR-61 (palivizumab), were cloned as Fab fragments with a C-terminal 3xMyc-1xHis8 tag, or as effectorless IgGs incorporating a human IgG1 Fc LALA-PG mutation [42], as previously described [18]. For controlled asymmetric heavy-chain pairing, we used knobs-into-holes technology in the context of a human IgG4 Fc as described by Carter [43], incorporating a stabilizing S228P mutation and mutations reducing immune effector function (F234A, L235A; FALA) as described by Dumet et al. [44]. T-cell engagers were equipped with an SP34-derived anti-CD3 binding domain (VR-43) [45] and cloned as bivalent antibodies in the 1+1 scFv-Fab-Fc format [45] or as trivalent antibodies in the 2+1 scDb-Fab-Fc format [46]. In bivalent 1+1 WiFi bodies, the anti-CD3-binding scFv arm was replaced with a VR-58 scFv, which was held constant across all TCRm antibody pairings. All constructs were produced in collaboration with Biointron (Beijing, China) as described previously [18]. Briefly, heavy and light chain sequences were synthesized and cloned into pcDNA3.4 expression vectors, followed by co-transfection into CHO-K1 cells. Proteins were expressed in suspension culture for 4-6 days at 37°C, 120 rpm, and 8% CO2. IgG antibodies were purified via protein A affinity chromatography followed by preparative size-exclusion chromatography. Fab fragments were purified using Ni-NTA resin under standard conditions. Protein purity, monomeric state, and integrity were assessed by analytical size-exclusion chromatography (SEC).

### Human cell lines

NALM-6 cells (ACC 128 from DMSZ) and T2 cells (ACC 598 from DMSZ) were grown in RPMI-1640 medium (Thermo Fisher Scientific). A375 cells (CRL-1619 from ATCC) were grown in DMEM medium with 4.5 g L-1 glucose (Thermo Fisher Scientific). Media were supplemented with 10% heat-inactivated fetal calf serum (FCS), and cells were cultured at 37°C and 5% CO2.

### Human Patient-derived Xenograft models

All PDX tumors were grown, harvested and characterized by EPO Berlin Buch GmbH. Tumors were preselected for positivity for HLA-A*02:01 as well as MAGEA4 as confirmed by RNAseq. Such tumors were purchased from EPO Berlin Buch GmbH.

### Immunopeptidomics-based characterization of endogenous MAGE-A4 pHLA complex presentation in PDX models and primary cell lines

First, cell lines were harvested and washed with PBS before freezing pellets for immunpeptidomic anlysis. The cells were lysed in 8 mL of 1% CHAPS (Millipore Sigma) supplemented with protease inhibitor cocktails (Thermo Scientific) for 1 hour at 4°C, after which the lysates were centrifuged at 20,000 g for 1 hour at 4°C, and the supernatant was collected. For PDX tumor samples frozen tumors were homogenized in CHAPS solution before centrifugation with identical conditions. For the immunopurification of HLA-I ligands, 0.5 mg of W6/32 antibody (Bio X Cell) was conjugated to 40 mg of CN Br-activated sepharose (Cytiva) and incubated with the protein lysate overnight. HLA complexes along with binding peptides were then eluted five times using 1% trifluoroacetic acid (TFA). These peptides and HLA-I complexes were separated using C18 columns (Sep-Pak C18 1 cc Vac Cartridge, 50 mg sorbent, 37-55 *µ*m particle size, Waters), which were pre-conditioned with 80% Acetonitrile (ACN) (Millipore Sigma) and equilibrated with three washes of 0.1% TFA. Samples were loaded onto the columns two times, washed two times with 0.1% TFA, and then eluted in 200 *µ*L of sequentially increased ACN concentrations - 15%, 30%, 40%, and 50% in 0.1% TFA. The four fractions were combined, dried using vacuum centrifugation, and stored at −80°C until further processing.

Afterwards, in-house C18 minicolumns were prepared using the following method: for the solid-phase extraction of a single sample, two small disks (1 mm diameter each) of C18 material were punched out from CDS Empore C18 disks (Thermo Fisher Scientific). These disks were then placed at the bottom of a 200 *µ*L Axygen pipette tip (Thermo Fisher Scientific). The columns were initially washed with 100 *µ*L of 80% ACN in 0.1% TFA and subsequently equilibrated three times with 100 *µ*L of 1% TFA. Fluids were passed through the column by centrifugation in a mini tabletop centrifuge, and the eluates were collected in Eppendorf tubes. Dried samples were then resuspended in 100 *µ*L of 1% TFA, loaded onto the columns, washed two times with 100 *µ*L of 1% TFA, and then run dry. Finally, the samples were eluted with 60 *µ*L of 60% ACN/0.1% TFA, and the resulting sample volume was further reduced by vacuum centrifugation.

Samples were analyzed using high-resolution and high-accuracy liquid chromatography-tandem MS on a Vanquish Neo ultra-high-performance liquid chromatography system coupled to an Orbitrap Exploris 480 mass spectrometer (Thermo Fisher Scientific). Peptides were separated on an in-house packed C18 analytical column with a 75 *µ*m inner diameter, 20 cm length, and 1.9 *µ*m beads (Dr Maisch Reprosil-Pur C18-AQ). The chromatographic gradient was applied at a flow rate of 250 nL/min, starting with 2% buffer B, which consisted of 90% acetonitrile and 0.1% formic acid, and 98% buffer A, which consisted of 3% acetonitrile and 0.1% formic acid. Over the first minute, the gradient increased to 5% buffer B, followed by a linear increase to 20% buffer B over the next 34 min. Subsequently, the gradient reached 30% buffer B over a 10 min period and further increased to 60% buffer B over the next 3 min. The system was then washed at 90% buffer B for 6 min before re-equilibration to 2% buffer B. MS was operated in data-dependent acquisition mode with a cycle time of 1 s. Full MS1 spectra were acquired at a resolution of 60,000 in a mass range of 300-1,600. Precursor ions were isolated with a window of 1.2 m/z and fragmented using a normalized collision energy of 28%. MS2 spectra were acquired at a resolution of 30,000 with an automatic gain control target of 100% and a maximum injection time of 150 ms. The dynamic exclusion time was set to 30 s and the intensity threshold specified to 10,000. Only precursor ions with charge states ranging from +1 to +4 were selected for fragmentation.

MS data were processed using Byonic software (V.2.7.84, Protein Metrics) and Peaks software (V.11, Bioinformatics Solutions) on a custom-built server equipped with 4 Intel Xeon E5-4620 8-core CPUs at 2.2 GHz and 512 GB of RAM (Exxact Corporation). The mass accuracy was set to 6 ppm for MS1 and 20 ppm for MS2. Digestion was specified as unspecific, allowing only precursors with charges of +1, +2, and +3 and a maximum mass of 2 kDa. Protein false discovery rate (FDR) was turned off to allow complete assessment of potential peptide identifications. Variable modifications included methionine oxidation; phosphorylation of serine, threonine, and tyrosine; and N-terminal acetylation. Samples were analyzed against a database containing UniProt Homo sapiens reviewed proteins, and common contaminants. Peptides ranging from 8 to 15 amino acids in length were selected with a minimum log probability value of 1.3, corresponding to p values of 0.05, and duplicates were removed. Ion intensities were exported from Byonic and Peaks for mirror plots. For peak area and retention time analysis, Skyline software (V.24.1, MacCoss Lab Software) was used. For quantitative mass spectrometry custom synthesized AQUA heavy peptides with ¿90% purity (Thermo Fisher) were spiked in at 30 fmol and molecules per cell calculated using cell numbers and comparison to the external standard.

To assign peptides that passed the MS quality filters to their most likely HLA complexes, we employed the NetMHCpan 4.1 algorithm (DTU Health Tech) with its default settings. Peptides with an affinity percentage rank below 2 were classified as binders.

### Detection of endogenous MAGE-A4 expression in primary cell lines by Western Blot analysis

1×10^7^ cells were washed with cold 1xPBS, lysed in RIPA buffer (ThermoFisher Scientific) supplemented with protease inhibitor cocktail (ThermoFisher Scientific), and incubated for 15 minutes on ice with gentle shaking. Lysates were cleared by centrifugation (14,000 x g, 15 minutes), and protein concentration was determined using the Pierce BCA Protein Assay Kit (ThermoFisher Scientific). 20 *µ*g total protein per sample was separated on a 12% Tris-Glycine gel (Novex WedgeWell, ThermoFisher Scientific) and transferred to a nitrocellulose membrane (ThermoFisher Scientific), using the iBright pre-stained protein ladder (ThermoFisher Scientific) as a molecular weight marker. Membranes were blocked in 5% milk powder (Carl Roth) in TBS-0.05% Tween20 (TBS/T) for 1-2 hours at room temperature, then incubated overnight at 4°C with primary antibodies: rabbit anti-human MAGE-A4 (Cell Signaling; 1:1000) and mouse anti-beta-tubulin (Invitrogen; 1:3000). After three 5-minute TBS-T washes, membranes were incubated with HRP-conjugated secondary antibodies (goat anti-rabbit and goat anti-mouse IgG H&L, Abcam; 1:2000 each) for 1 h at room temperature. Following three further washes, signals were developed with ECL substrate (SuperSignal West Pico PLUS, ThermoFisher Scientific) and detected on an iBright FL1500 imaging system.

### Peptide-HLA array production and HighSCORE measurements

Peptide-HLA array production and HighSCORE measurements were performed essentially as previously described [21]. Peptides were purchased from Peptides & Elephants GmbH (Hennigsdorf, Germany), and biotinylated, stabilized HLA-A*02:01 variant was used. VR-4 and VR-6 were measured in Fab format to avoid avidity effects. Binding curves were fitted and analyzed using an in-house version of Anabel and a 1:1 Langmuir binding model [47]. Each unique peptide-HLA combination was measured in 16 replicates to reduce the overall false-positive rate.

### T2 cell peptide loading and pHLA surface display

Peptide loading of T2 cells was performed essentially as previously described [18]. In brief, T2 cells and peptide solution were mixed directly in 96-well plates or pre-mixed in Falcon tubes using AIM-V medium (Thermo Fisher Scientific); both approaches yielded equivalent peptide loading. Peptide loading was performed at 37°C and 5% CO2 for 17-18h. Following incubation, T2 cells were washed and immediately subjected to subsequent assays.

### Flow cytometry-based binding assays

TCRm binding analysis was performed using a full 8-point dose titration (starting with 250 nM to 0.02 nM with 1:4 dilution). Briefly, peptide-loaded T2 cells were incubated with TCRm antibodies (VR-4, VR-6, and VR-58) in IgG format. After 1 h of incubation, the cells were stained with a secondary FITC-labeled anti-human Fc detection antibody (Thermo Fisher Scientific, #A18818). In addition, the cells were stained with LIVE/DEAD™ Fixable Violet Staining Solution (Thermo Fisher Scientific, #L34964) to distinguish between viable and dead cells. For peptide loading control, T2 cells were stained with an AF647 anti-B2M antibody (Thermo Fisher Scientific, #MA5-18119). Flow cytometry was performed using an Attune NxT flow cytometer (Thermo Fisher Scientific).

### Primary human T-cell Generation and Expansion

Cryopreserved peripheral blood mononuclear cells (PBMCs) from healthy donors (Biomol GmbH) were thawed and resuspended in RPMI-1640 medium (Thermo Fisher Scientific) supplemented with 10% heat-inactivated fetal calf serum (FCS) and 500 U/ml recombinant human IL-2 (PeproTech, #200-02). PBMCs were activated with plate-bound anti-human CD3 (UCHT1,BioLegend, #300402) and anti-human CD28 antibodies (clone CD28.2, BioLegend, #302902), 1 *µ*g/ml each. After 48-72 h, ¿99% of the cells were T-cells as evidenced by anti-human CD3 staining. Primary human T-cells were maintained in RPMI-1640 medium supplemented with 10% FCS and 100 U/ml recombinant human IL-2 until they were used in subsequent assays.

### T-cell-mediated cytotoxicity and T-cell activation assays

T-cell-mediated cytotoxicity was measured using flow cytometry-based assay. Briefly, target cells, either peptide-loaded T2 cells or cancer cells, were labeled with Cell-Trace™ Far Red (Thermo Fisher Scientific, #C34572) and incubated for 4h and 24h at 37°C and 5% CO2 with primary human T-cells at a 1:1 Effector to target (E:T) ratio in the presents of TCEs in 96-well plates. Target cells incubated without TCEs and primary human T-cells served as controls for spontaneous cell death during the assay. After 4h and 24h, cells were harvested, stained with LIVE/DEAD™ Fixable Violet staining solution (Thermo Fisher Scientific #L34964) and analyzed by flow cytometry (Attune NxT, Thermo Fisher Scientific). CellTrace™ Far Red positive target cells were gated, and the percentage of dead target cells (LIVE/DEAD™ Fixable Violet positive cells) was determined. Specific killing was calculated using the following formula: % specific lysis = [(% target cell death in treated sample - % spontaneous cell death target cells) / (100 - % spontaneous cell death target cells)] *100.

In addition, T-cell activation following target cell encounter was assessed by measuring the upregulation of CD69 and CD25 and IFN-γ release. Briefly, primary human T-cells were stained with APC-eFluor™ 780 - labeled anti-human CD69 and PE-eFluor™ 610-labeled anti-human CD25 (eBioscience, #47-0699-42 and #61-0259-42). In addition, T-cells were stained with FITC-labeled anti-human CD8 (RPA-T8) and PerCP-Cyanine5.5-labeled anti-human CD4 (RPA-T4) antibodies (eBioscience #11-0088-41 and #45-0049-42). To determine IFN-γ release after 24h incubation, supernatants were collected and cytokine levels were determined using a commercial ELISA Kit (Invitrogen #88-7316-77). T-cell-mediated cytotoxicity assay and T-cell activation assay were performed simultaneously.

### Structural determination of TCRm-pHLA complexes by cryo-EM

#### Production and purification of HLA-A*02:01 and hβ2m inclusion bodies

MHC class I heavy chain (HLA-A*02:01 Y84C/A139C variant) and human β2-microglobulin (hβ2m) were produced separately as inclusion bodies in E. coli strain BL21(DE3), using a protocol modified from previously described procedures [48]. Cells were harvested by centrifugation, re-suspended in lysis buffer (50 mM Tris pH 8.0, 25% (w/v) sucrose, 1 mM EDTA, 10 mM DTT, 1 mM PMSF), and lysed by sonication. Inclusion bodies were pelleted by centrifugation (22,000 x g, 20 min, 4 °C) and sequentially washed by cycles of resuspension, sonication and centrifugation in detergent buffer (50 mM Tris pH 8.0, 25% (w/v) sucrose, 1% (v/v) Tween20, 5 mM EDTA, 2 mM DTT), twice in urea/salt buffer (50 mM Tris pH 8.0, 2 M NaCl, 2 M urea, 2 mM DTT), and in Tris-buffer (50 mM Tris pH 7.5, 150 mM NaCl, 0.5 mM PMSF). Washed inclusion bodies were solubilized for 48 h at 4 °C in denaturation buffer (50 mM HEPES pH 6.5, 6 M guanidinium hydrochloride, 0.5 mM PMSF, 100 *µ*M β-mercaptoethanol) with gentle agitation, and insoluble material was removed by two successive rounds of centrifugation (22,000 x g, 20 min, 4 °C). The concentration of the soluble denatured protein was determined by UV absorbance at 280 nm using sequence-derived extinction coefficients, and aliquots were stored at −80 °C until use.

#### Refolding and purification of peptide-receptive HLA-A*02:01 complexes

Peptide-receptive HLA-A*02:01(Y84C/A139C)-hβ2m complexes were generated by dilution refolding in the presence of the dipeptide glycyl-L-methionine (GM), using a protocol modified from previously described methods [49, 50]. In brief, denatured hβ2m was slowly added to cold refolding buffer containing 100 mM Tris-HCl, pH 8.0, 500 mM L-arginine hydrochloride, 2 mM EDTA, 5 mM reduced glutathione, 0.5 mM oxidized glutathione, 100 *µ*M PMSF, and 10 mM GM to a final hβ2m concentration of 2 *µ*M. The mixture was stirred for 1 h at 4 °C. Denatured HLA-A*02:01(Y84C/A139C) heavy chain was subsequently added in portions to a final concentration of 1 *µ*M. The refolding reaction was incubated for 4 days at 4 °C with continuous gentle stirring. The refolding mixture was clarified by filtration through a 0.22 *µ*m membrane and concentrated approximately 50-fold by ultrafiltration using a 30 kDa cutoff membrane. The concentrated refolding mixture was purified by size-exclusion chromatography (SEC) using a HiLoad 16/600 Superdex 200 pg column equilibrated in SEC buffer containing 50 mM Tris-HCl, pH 8.0, 150 mM NaCl, and 2 mM GM. Fractions corresponding to the expected HLA-A*02:01(Y84C/A139C)-hβ2m heterodimer were analyzed by SDS-PAGE and pooled for subsequent peptide loading.

#### Assembly and purification of pHLA-Fab complexes for cryo-EM

Peptide-receptive HLA-A*02:01(Y84C/A139C)-hβ2m complex was incubated at a final concentration of 14 *µ*M with 140 *µ*M MAGE-A4 peptide (GVYDGREHTV), corresponding to a 10-fold molar excess of peptide, for 15 min on ice. For each complex preparation, VR-4, VR-6, or VR-58 Fab antibody was subsequently added at an equimolar concentration relative to the pHLA complex, and the mixture was incubated for an additional 30 min on ice. The resulting pHLA-Fab complex was purified by SEC using a Superdex 200 Increase 10/300 GL column equilibrated in Dulbecco’s phosphate-buffered saline (DPBS). Fractions corresponding to the pHLA-Fab complex were identified based on the chromatographic elution profile and analyzed by SDS-PAGE. Appropriate fractions were pooled, and the protein concentration was determined by measuring the absorbance at 280 nm using a sequence-derived extinction coefficient calculated for the complete pHLA-Fab complex. The sample was concentrated to approximately 2.0 mg/mL by centrifugal ultrafiltration using a 10 kDa cutoff Amicon Ultra filter device. Anti-human β2-microglobulin Fab BBM.1 [51] was added to the purified pHLA-Fab complex at an equimolar ratio relative to the pHLA complex. The mixture was incubated for 30 min on ice and subsequently subjected to a second SEC step using a Superdex 200 Increase 10/300 GL column equilibrated in DPBS. Fractions corresponding to the assembled pHLA-VR Fab-anti-hβ2m Fab complex were analyzed by SDS-PAGE and pooled. The purified complex was concentrated to approximately 2.5 mg/mL by centrifugal ultrafiltration using a 10 kDa cutoff Amicon Ultra filter device and maintained on ice until preparation of cryo-EM grids.

#### Cryo-EM sample preparation

Protein samples were diluted to 0.5 mg/mL for cryo-EM grid preparation. Protein samples were diluted to 0.5 mg/mL for cryo-EM grid preparation. Quantifoil R1.2/1.3 300-mesh copper grids were washed with chloroform and glow-discharged at 15 mA for 90 s using a PELCO easiGlow device. A 4 *µ*L aliquot of sample was applied to each glow-discharged grid, blotted for 4 s at a nominal blot force of 20, and plunge-frozen in liquid ethane using a Vitrobot Mark IV (Thermo Fisher Scientific).

#### Cryo-EM data collection

All cryo-EM datasets were recorded in energy-filtered transmission electron microscopy mode using a Titan Krios G4 microscope (Thermo Scientific) operated at 300 kV. Electron-optical alignment was performed with EPU 3.15 (Thermo Scientific). Images were recorded in electron-counting mode using a Falcon 4i direct electron detector (Thermo Scientific) with a nominal magnification of 215,000x, corresponding to a calibrated pixel size of 0.573 Å, and with a nominal defocus between −1.0 and −2.0 *µ*m. Dose fractionated movies were recorded in in electron-event representation format at an electron dose rate of about 8 e-/pixel/s for 2.2 s, corresponding to a total dose of approximately 50 e-/Å 2.

#### Cryo-EM image processing

All cryo-EM datasets were processed using the same approach in CryoSPARC 4.7.1 [52]: Beam-induced motion correction, dose-weighted image generation, and contrast transfer function (CTF) parameter estimation were done using patch motion correction and patch CTF estimation. Images with estimated resolution over 5.0 Å, and astigmatism over 400 Å were discarded. Particles were picked with blob picker and TOPAZ [53] for further processing. Two-dimensional classification, initial model generation, heterogeneous refinement, homogeneous refinement, non-uniform refinement were done in CryoSPARC [52, 54]. CTF refinement and Bayesian polishing were performed using CryoSPARC’s implementation [55, 56]. The pixel size of the particles for final reconstruction was 0.8022 Å. Fourier shell correlation (FSC) curves and local resolution estimation were done in CryoSPARC for all final maps. Local resolution estimation, Fourier shell correlation (FSC) curves, angular distribution of particles, cryo-EM densities around MAGE-A4 peptide, and cryo-EM map-to-model fitting FSCs of the final cryo-EM maps are shown in Fig. S9. Further, the chart of cryo-EM data processing is shown in Fig. S10. A summary of map qualities is shown in Table S4.

#### Model building and geometry refinement

The first atomic model was built into the cryo-EM density map of the pHLA-VR6-BBM.1 complex using ISOLDE within ChimeraX 1.11 and Coot 0.9.8.92 [56–59], based on the crystal structures of HLA-MAGE-A4 (PDB 8FJA) [23] and BBM.1 (PDB 9UVI) [51], and the VR6 model generated by AlphaFold [60], followed by real space refinement in Phenix 1.20.1 [61, 62]. The refined model was manually inspected and adjusted if necessary. The model of pHLA-VR6-BBM.1 complex was used as a template to build the other atomic models. All the models were validated by MolProbity implemented in Phenix [63]. Map-to-model cross-validation was performed in Phenix [61]. Comprehensive information on the cryo-EM data collection, refinement, and validation statistics is shown in Table S3. Molecular Operating Environment (MOE), version 2024.0601 (Chemical Computing Group ULC, Montreal, QC, Canada), was used to calculate interaction energies and to visualize the structures shown in this study. ChimeraX was used for cryo-EM map visualization.

### EpiTox-based peptide-centric off-target liability prediction and annotation

EpiTox, a multi-modular computational platform for proteome-wide off-target prediction, was used to define the off-target landscape for both the MAGE-A4 KVL nonapeptide and the MAGE-A4 GVY decapeptide, as detailed in Figure 2 A. In brief, EpiTox’s performed a proteome-wide sequence similarity search for each target peptide, and candidates were ranked using complementary scores: a Bi-feature score incorporating predicted HLA-A*02 binding affinity and a Multi-feature score based on the peptide’s mismatch pattern relative to the target (anchor positions, backbone, or both) and biophysical features, as detailed in [20]. Tissue-specific expression relevance was subsequently incorporated via a decision matrix to prioritize candidate off-target peptides for lab-based experimental validation [20]. For the MAGE-A4 GVY off-target landscape, this approach was used to select the labeled training dataset, which was combined with the complete positional XScanning variant panel of the MAGE-A4 wild-type peptide for HighSCORE-based kinetic characterization and subsequent EpiPredict model training [18]. EpiTox was also used to annotate the peptides predicted by EpiPredict and analyzed in this study.

Off-target pools for both peptide targets (GVYDGREHTV and KVLEHVVRV) were generated independently using EpiTox. For each peptide, EpiTox estimates three scores, Bi-feature, Multi-feature, and experimental-evidence, capturing complementary aspects of off-target profile and physiological relevance. Gene expression data (nTPM values obtained from the Human Protein Atlas [41] were used to classify off-targets into five expression tiers, based on two dimensions: expression breadth (the number of tissues, out of 40 profiled, in which a gene is expressed) and critical-tissue expression (expression level in a predefined set of critical tissues: heart muscle, liver, skeletal muscle, kidney, lung, cerebral cortex, spleen, thymus, and lymph node). The five tiers, in order of decreasing concern, are: Broad and Critical, Broad and Non-critical, Critical only, Restricted, and Low/Absent. Tiers were assigned hierarchically, such that each tier captures only the genes not already assigned to a tier above it. Broad and Critical: expressed in ≥20 tissues, with expression ¿10 nTPM in at least one critical tissue; Broad and Non-critical: expressed in ≥20 tissues, without exceeding 10 nTPM in any critical tissue; Critical only: expression exceeding 10 nTPM in at least one critical tissue; Restricted: expressed in 5-19 tissues; Low/Absent: expressed in fewer than 5 tissues.

Tier thresholds (≥20 tissues for broad expression; ¿10 nTPM for critical-tissue expression) were derived from the empirical distributions of expression breadth and critical-tissue expression across both pools.

The risk core was defined for each target as the subset of off-targets satisfying all three of the following criteria simultaneously: (i) Bi-feature score above the target-specific median, (ii) Multi-feature score above the target-specific median, and (iii) expression tier classified as Broad and Critical or Critical only. This convergent filter identifies off-targets that cannot be dismissed on any single axis. Core sizes were compared between the two targets using a two-proportion z-test.

### EpiPredict-based proteome-wide TCRm-centric off-target prediction

Proteome-wide off-target risk assessment was performed using EpiPredict, a TCRm-specific machine learning framework trained on high-throughput kinetic binding data, as previously described [18]. In brief, antibody-specific neural network models were trained on labeled kinetic interaction data generated by HighSCORE screening against a peptide panel comprising positional X-scan variants of the MAGE-A4 GVY decapeptide and EpiTox-predicted sequence-similar off-target peptides. This approach was applied to VR-4 and VR-6 as previously described [18], and independently extended to VR-58 using the same peptide panel and screening methodology. The trained models were applied to a library of proteome-derived decapeptides predicted to be presented by HLA-A*02 (NetMHCpan [64] rank ≤2), generating a TCRm-specific binding probability HighSCORE for each peptide. Peptides exceeding a probability threshold of 50% were prioritized as candidate off-targets for experimental validation via HighSCORE and T2 cell-based binding assays.

### ParaPredict: structural modeling with Chai-1 and MOE-based refinement

Structural modeling of TCRm-pHLA complexes was performed using Chai-1 [65] in single-sequence mode, with antibody-specific restraints derived from CDR-peptide/HLA contacts found in PDB-structures and from wet-lab data, followed by MOE-based relaxation and reference-free interface scoring (buried surface area, clash density, hydrogen bonding, and ensemble stability) to rank and validate candidate models [18].

### Labeled kinetic dataset generation

The labeled kinetic training dataset used for EpiPredict model development was generated as previously described [18]. Briefly, EpiTox-based in silico off-target prediction identified candidate peptides with up to five substitutions relative to the MAGE-A4 GVY epitope, filtered for predicted HLA-A*02:01 presentation and expression relevance. These were combined with a complete single-amino-acid substitutional scan (XScanning) of the wild-type peptide, yielding 530 unique peptides. Binding interactions with VR-4, VR-6, VR-57 and VR-58 were quantified using HighSCORE [21] and analyzed by Anabel software [47], and peptides were classified as binders, non-binders, or ambiguous based on binding curve validity, replication, and quality control criteria [18]. The resulting labeled datasets (530 peptides per antibody) served as direct input for EpiPredict model training.

### Unlabeled 10-mer peptide dataset generation

The unlabeled 10-mer peptide space used for proteome-wide inference was generated as previously described [18]. Briefly, all canonical and isoform sequences from UniProt (release February 2025; 34,184 protein sequences) were parsed into a sliding-window library of ¿10 million unique 10-mers, filtered for predicted HLA-A*02:01 presentation using NetMHCpan 4.1 (rank threshold ≤2), yielding a final set of 316,855 unique peptides used as the unlabeled prediction space.

### Databases

Our antibodies and molecules are fully tracked using the Genedata Biologics (GDB) system [66].

## ACKNOWLEDGMENTS

The authors gratefully acknowledge Quinton Holland for his continuous support and his constructive drive in pushing the boundaries of what we believed achievable. The authors thank the Central Electron Microscopy Facility at Max Planck Institute of Biophysics for technical support and access to instrumentation. AI use: GPT-5.5 and Claude Sonnet 4.5 and Sonnet 5 models were employed exclusively for language refinement during manuscript preparation.

## AUTHOR CONTRIBUTIONS

Conceptualization (S.S, J.B., M.G.K); Methodology (S.S., H.A.-H., S.K., A.S., P.H., S.L., G.R., C.R., T.-H.W. O.S., M.G.K.); Investigation (J.S., F.B., T.-H.W., T.H., S.Ö., M.S., T.G., L.S., N.K., G.J., J.Sc., P.M., J.G.); Data curation and Software (S.S., H.A.-H., S.K., A.S., B.L., O.S.); Structural models and analysis (T.-H.W, C.R., H.M.); Visualization (H.A.-H., S.K., O.S.); Supervision (S.S, O.S., C.R., F.A.H., K.H., B.S.); Writing-original draft (J.B.); Writing-review & editing (All authors).

## DATA, CODE, AND MATERIALS AVAILABILITY

All data supporting the findings of this study are included in the paper and its Supplementary Materials. Atomic coordinates and cryo-EM maps for the VR-4-, VR-6-, and VR-58-MAGE-A4/HLA-A*02:01 Fab complexes have been deposited in the Protein Data Bank and Electron Microscopy Data Bank; accession codes are not included in this preprint and will be provided with the peer-reviewed publication.

## Supplementary Material

**Figure S1.**
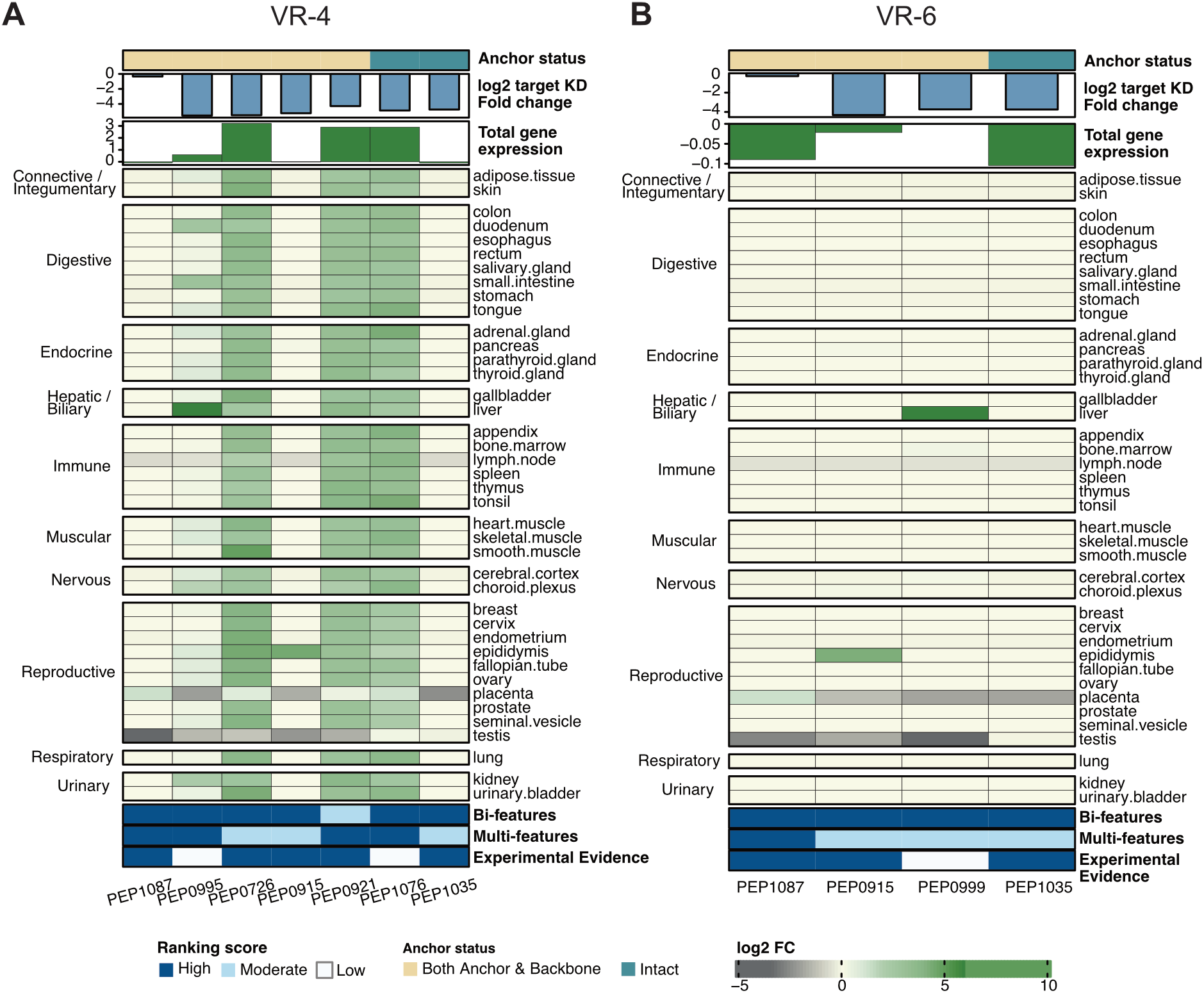
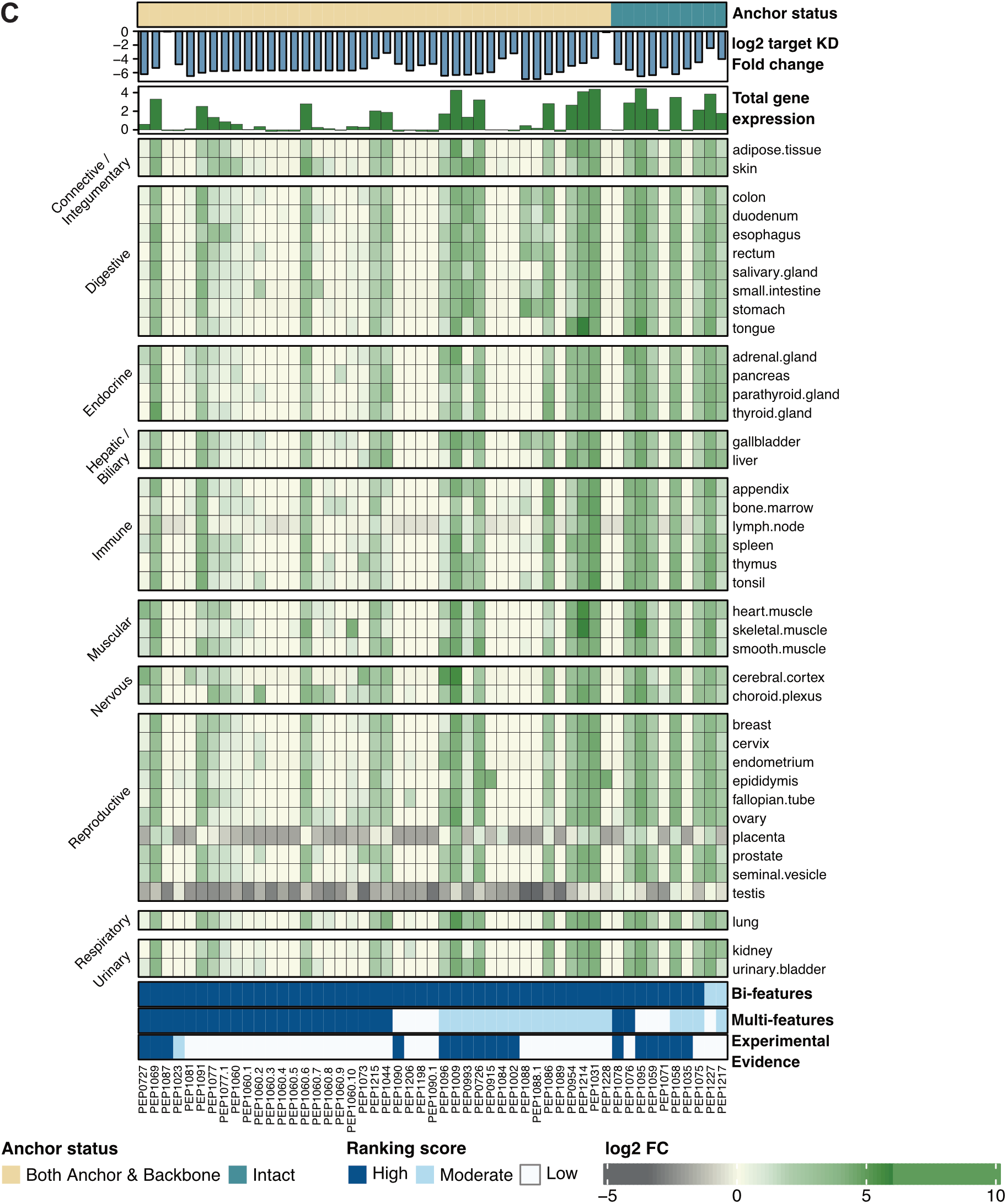
Annotated heatmap of confirmed EpiTox-predicted off-targets including tissue distribution and risk features. Confirmed binders of EpiTox-predicted sequence-similar peptides identified by HighSCORE-based screening for VR-4, VR-6, and VR-58. Heatmap showing gene expression profiles across multiple tissues for peptides confirmed as binders to **(A)** VR-4, **(B)** VR-6, or **(C)**) VR-58 in HighSCORE screening. Gene expression values are displayed as log2 fold-change relative to target tissue expression. Annotations, from top to bottom: anchor status, where “intact” indicates that all conserved residues at predicted HLA anchor positions are preserved (blue), and “Both Anchor and Backbone” indicates non-permissive amino acid substitutions at both anchor and backbone positions (beige); peptides carrying mismatches exclusively at anchor positions are not represented. Anchor positions refer to key peptide residues, typically at positions 2 and 10 for 10-mers like GVY, that make direct contacts with the HLA binding groove and are critical for stable pHLA complex formation; log2-transformed KD fold-change relative to the target peptide, where negative values indicate lower binding affinity than the target and positive values indicate higher affinity; bar plot showing total gene expression summed across all tissues per peptide. Bi-feature rank, Multi-feature rank, and Experimental Evidence score as described in Al-Hasani et al., 2025 [20]. Briefly, darker colors indicate higher risk of representing a physiologically relevant off-target based on sequence similarity and HLA-A*02 binding affinity (Bi-feature score), biophysical property similarity (Multi-feature score), and experimental evidence for the existence of the respective pHLA complex in humans (Experimental Evidence score), with dark blue across all three categories representing the highest risk. Orange boxes indicate peptides PEP-0726 and PEP-0727 selected for cell-based binding and cytotoxicity studies (see “TCRm-mediated cellular cytotoxicity”). PEP-0915 is MAGE-A11-derived. Peptide IDs appearing multiple times in the heatmap indicate that the corresponding sequence maps to more than one gene.

**Figure S2.**
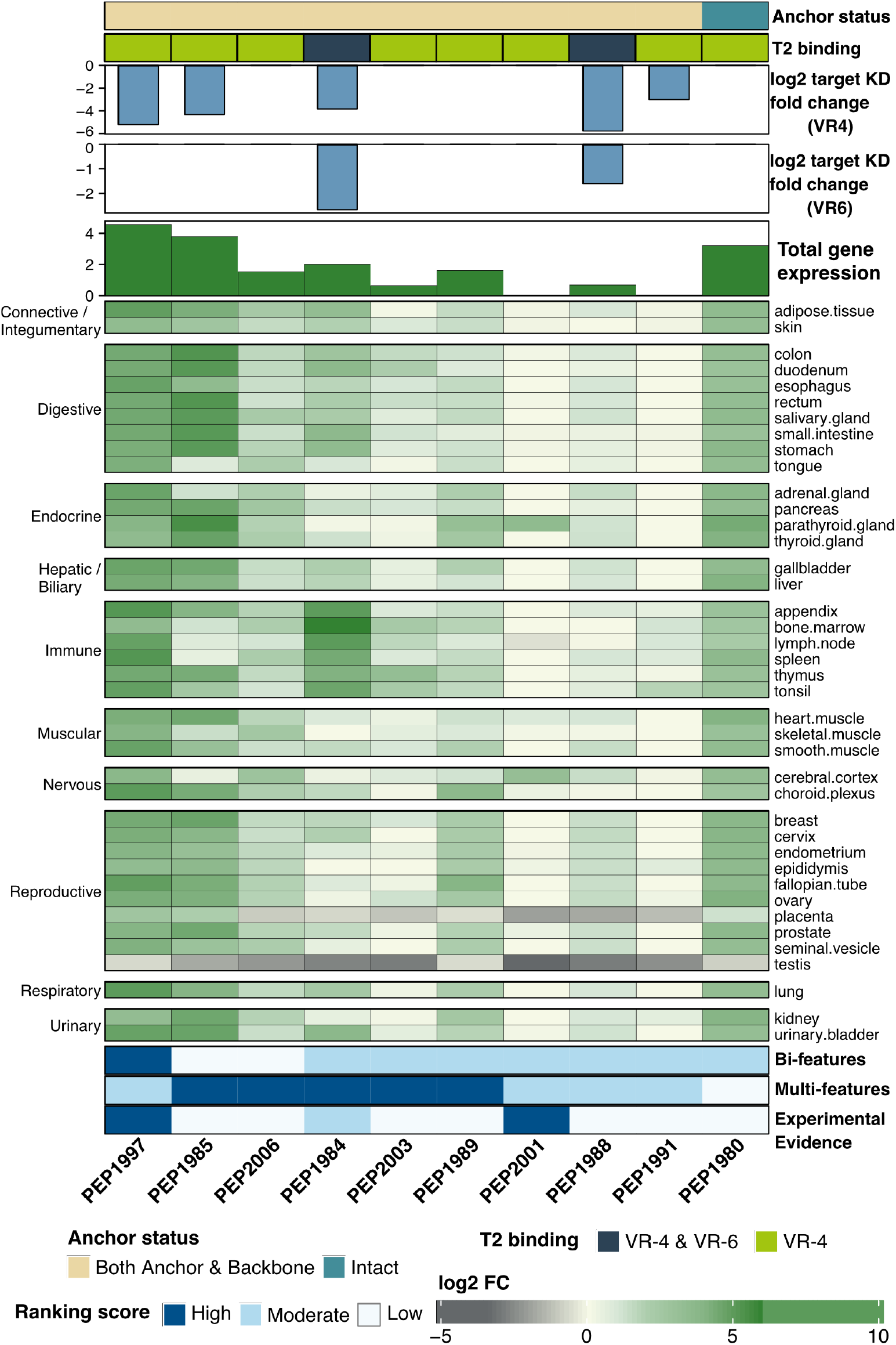
Annotated heatmap of confirmed EpiPredict-predicted off-targets including tissue distribution and risk features. Confirmed binders among EpiPredict-predicted sequence-similar peptides identified by HighSCORE-based screening for VR-4 and VR-6. Annotations are as described for EpiTox-predicted binders in Figure S1. Since full dose-response binding could not be achieved for all binders in the monovalent Fab format by HighSCORE, binding was verified at least qualitatively using T2 cell-based binding assays with bivalent IgGs (Figure 4 B; [18]).

**Figure S3.**
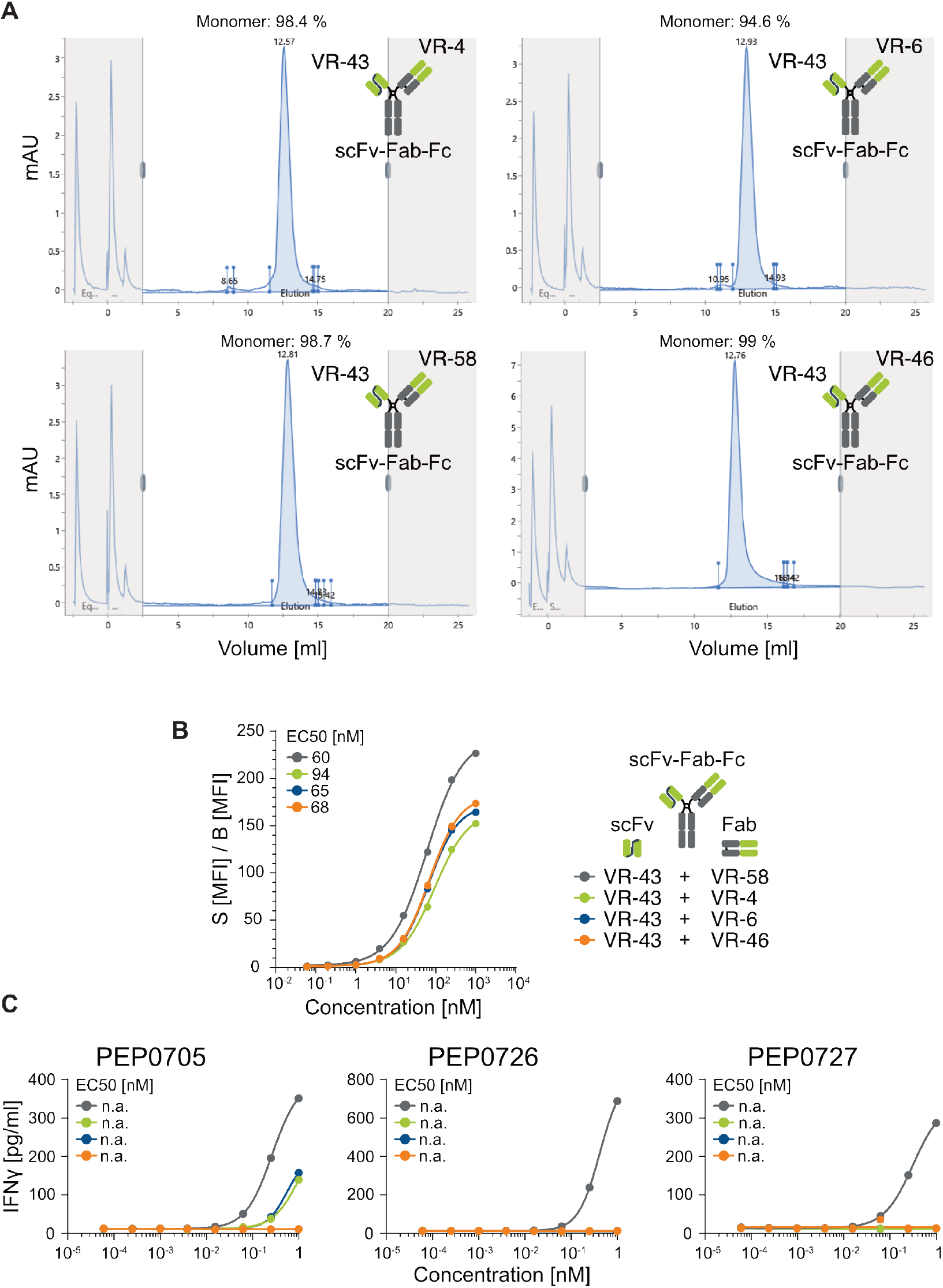
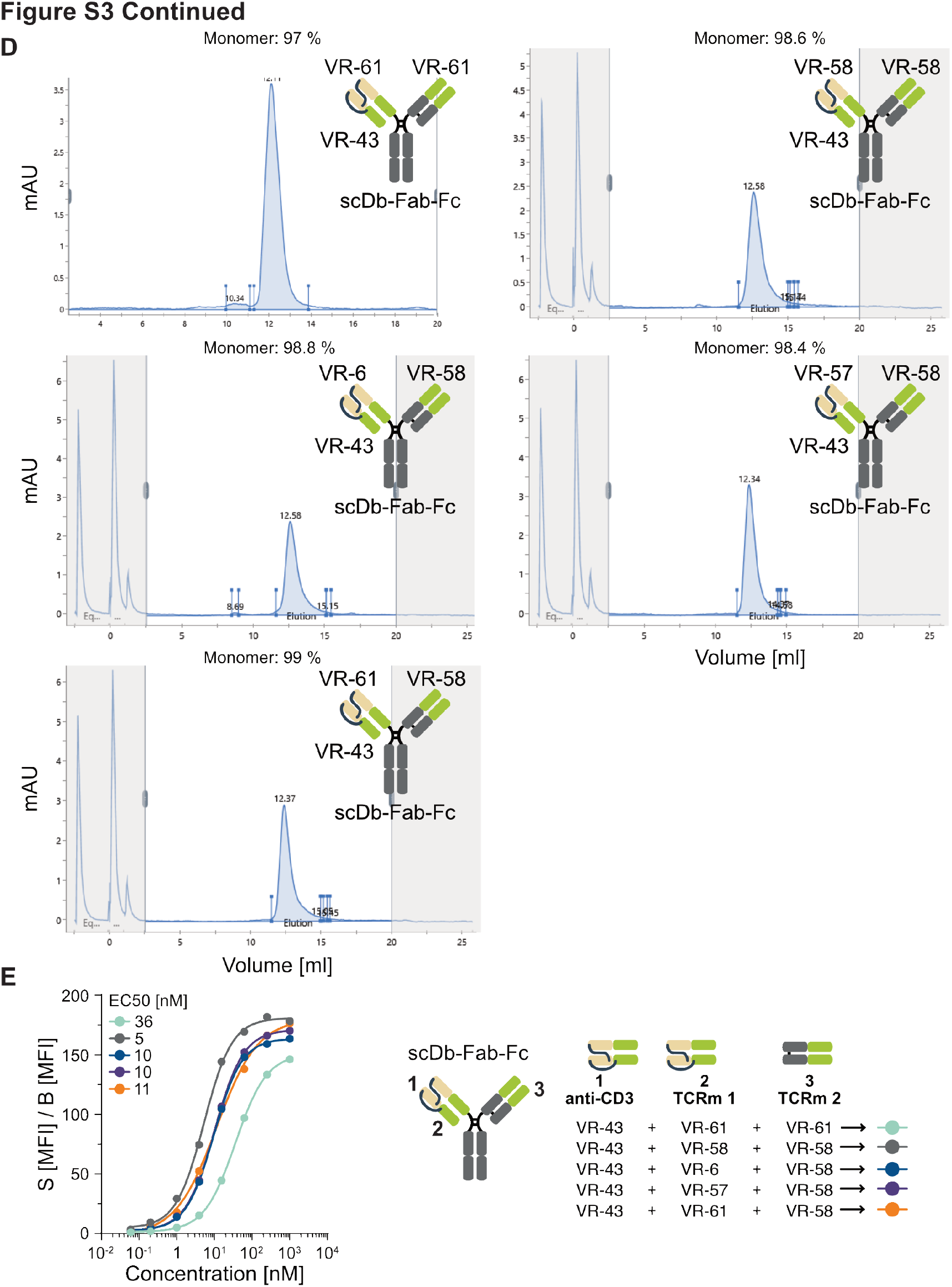
Characterization of bi- and tri-valent T-cell engagers. **(A)** Purity of bivalent scFv-Fab-Fc 1+1 T-cell engagers. TCEs were purified by protein A affinity chromatography followed by preparative SEC to at least 95% monomer content, as confirmed by analytical SEC; SEC profiles and monomer content [%] are indicated. **(B)** EC_50_ dose-response curves and values for binding of scFv-Fab-Fc 1+1 T-cell engagers to primary T-cells, mediated by anti-CD3 binding. **(C)** Bivalent scFv-Fab-Fc T-cell engagers containing different VR (variable region) combinations induced interferon-γ release from primary T-cells after 24 h of incubation with peptide-loaded T2 cells. **(D)** Purity of trivalent scDb-Fab-Fc 2+1 T-cell engagers. TCEs were purified as in **(A)** to at least 95% monomer content; SEC profiles and monomer content [%] are indicated. The position of the anti-CD3-binding domain (VR-43) within the scDb arm is indicated in brown. **(E)** EC_50_ dose-response curves and values for binding of trivalent scDb-Fab-Fc 2+1 T-cell engagers containing different VR combinations to primary T-cells, mediated by anti-CD3 binding. Schematic representation of the bivalent scFv-Fab-Fc 1+1 format **(A-C)** or trivalent scDb-Fab-Fc 2+1 T-cell engagers **(D, E)** indicating SP34-derived anti-CD3 VR-43 (brown) on the scFv or scDb arm and other variable binding moieties on the Fab arm **(A-C)** and scDb and Fab arm **(D, E)**.

**Figure S4.**
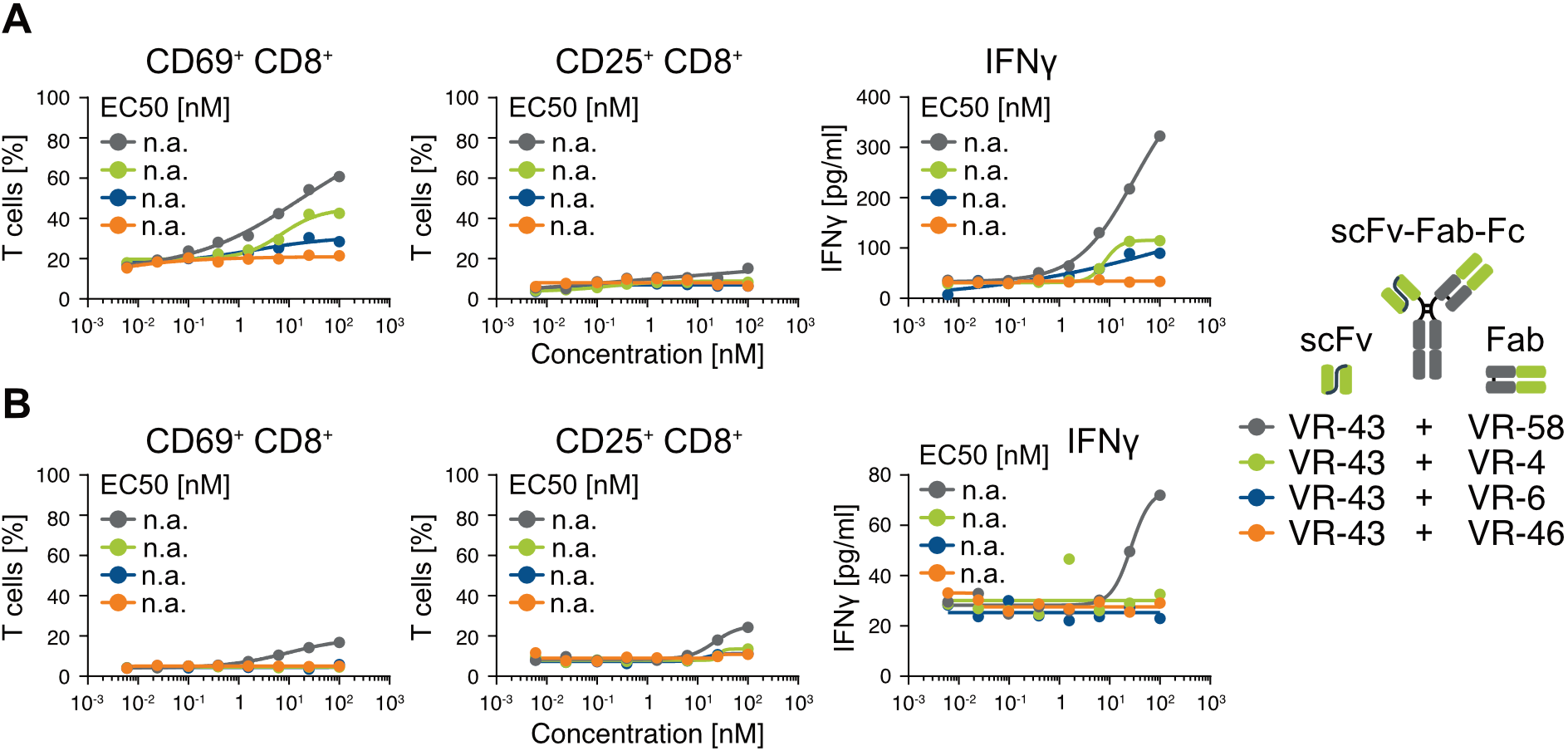
T-cell activation by bivalent T-cell engagers. **(A, B)** Activation of primary T-cells following incubation with A375 **(B)** or NALM-6 **(C)** target cells in the presence of scFv-Fab-Fc-based bivalent T-cell engagers, as evidenced by CD69 positivity after 4 h and CD25 positivity after 24 h by flow cytometry, and interferon-γ release after 24 h by ELISA. Schematic representation of the bivalent scFv-Fab-Fc 1+1 format, indicating SP34-derived anti-CD3 VR-43 on the scFv arm and other variable binding moieties on the Fab arm. n.a., no binding signal or binding signal insufficient to complete dose-response titration.

**Figure S5.**
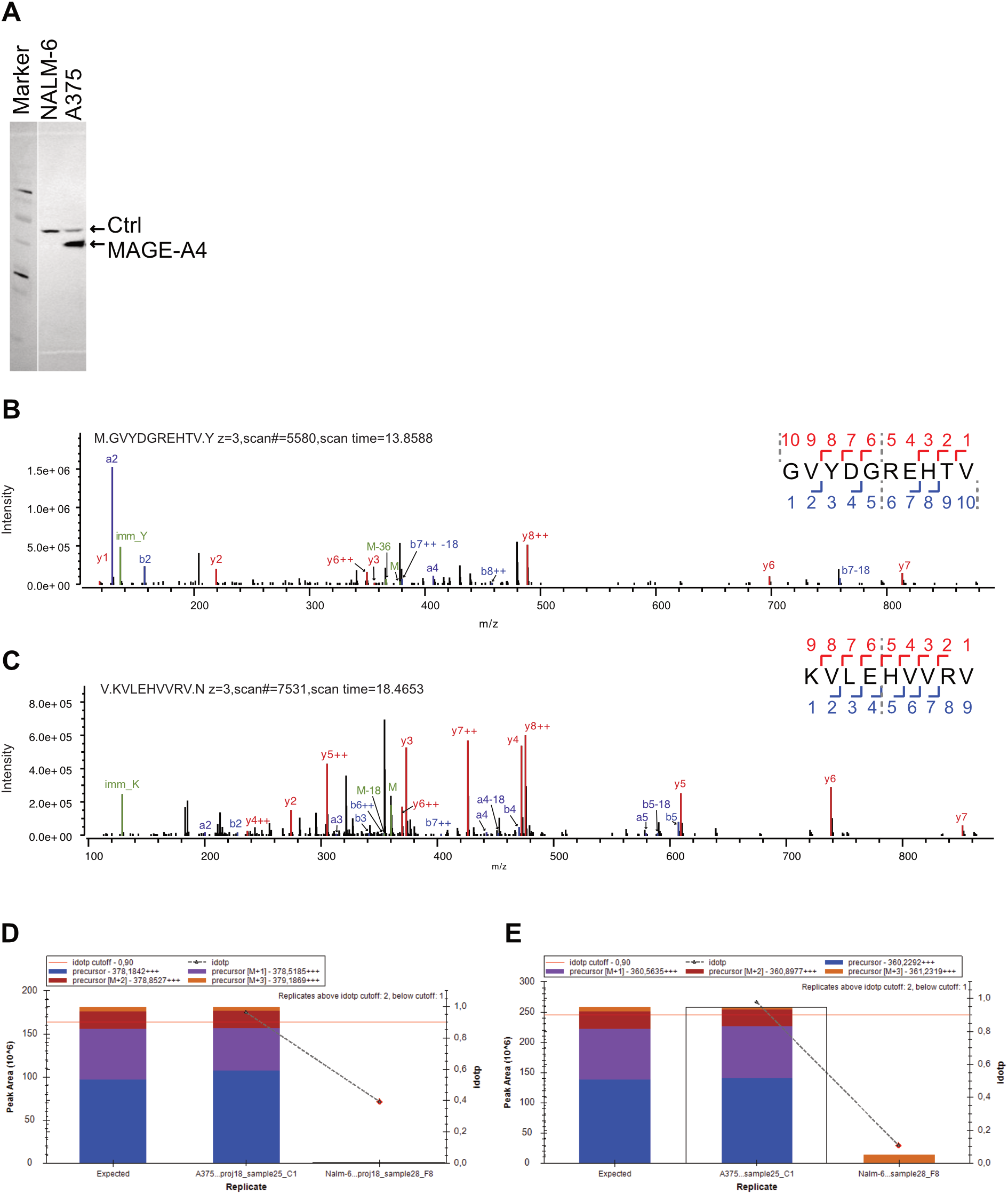
MAGE-A4 expression and peptide spectra and peak intensities of GVY and KVL peptides in A375 and Nalm6 cells. **(A)** Expression of MAGE-A4 in A375 and NALM-6 cell lines by Western blotting **(B, C)** Peptide spectrum matches generated by Byonic from GVY and KVL peptides. **(D, E).** Peak areas of the peptides detected in A,B as measured with Skyline. D shows detection of GVY peptide in A375 but not Nalm6 cells and E shows detection of KVL peptide in A375 but not Nalm6 cells.

**Figure S6.**
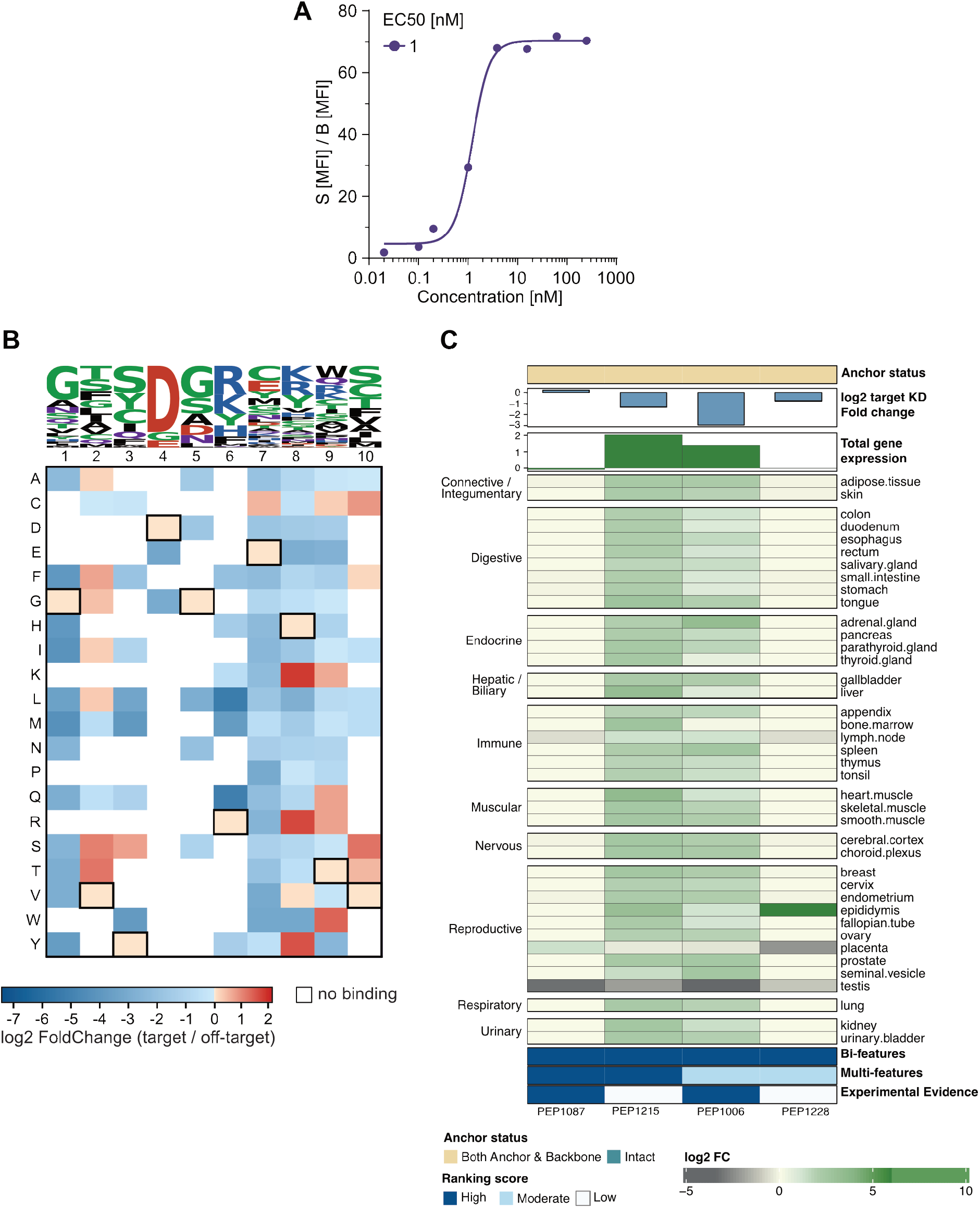
Characterization of the anti-MAGE-A4 binder VR-57. **(A)** EC_50_ binding titration of VR-57 against MAGE-A4 target peptide PEP-0705-loaded T2 cells (5 *µ*M loading concentration). **(B)** HighSCORE-based XScanning profile and logo plot, as described for Figure 3. **(C)** Annotated off-targets identified for VR-57 by HighSCORE-based off-target screening, using the same off-target panel as used for VR-4, VR-6, and VR-58.

**Figure S7.**
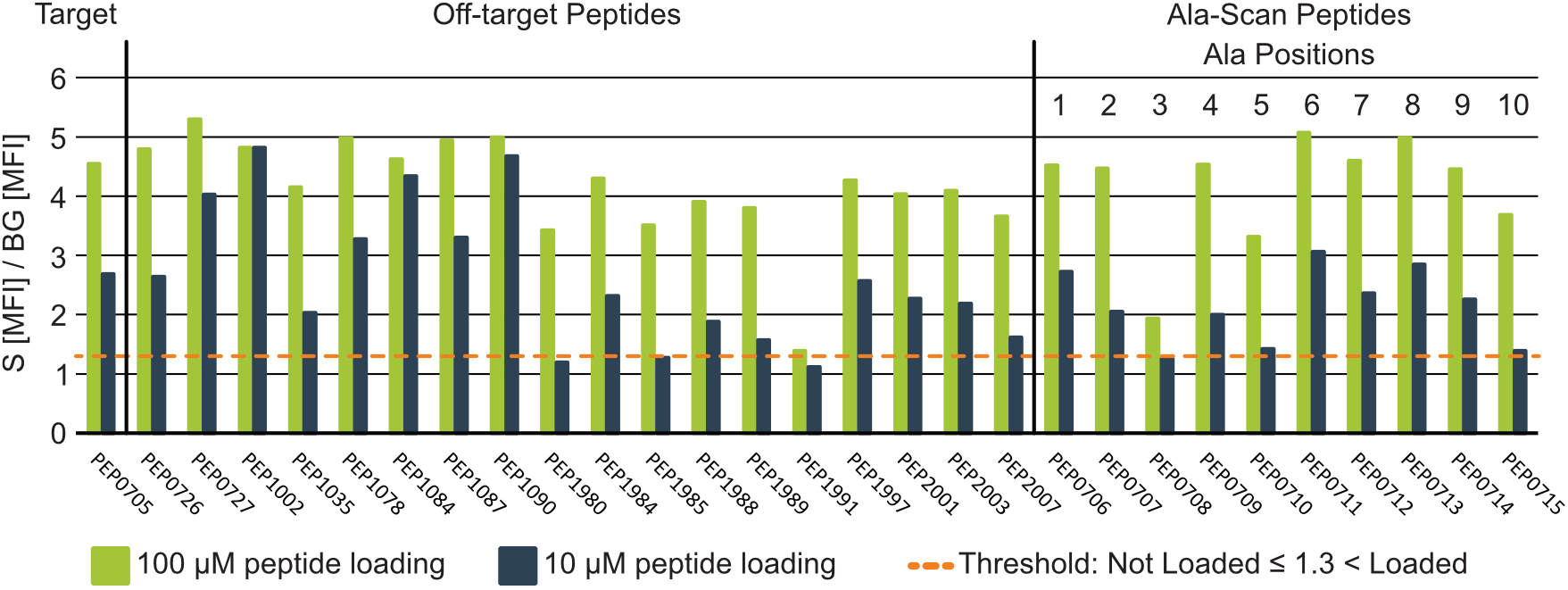
T2 cell peptide-loading validation. T2 cells were loaded with 100 µM or 5 µM of the wild-type decapeptide (PEP-0705), Ala-scan variants (PEP-0706-PEP-0715), off-target peptides predicted by EpiTox (PEP-0726, PEP-0727), or off-target peptides predicted by EpiPredict (PEP-1002, PEP-1035, PEP-1078, PEP-1084, PEP-1087, PEP-1090, PEP-1980, PEP-1984, PEP-1985, PEP-1988, PEP-1989, PEP-1991, PEP-1997, PEP-2001, PEP-2003, PEP-2007). To control for false negatives due to insufficient peptide presentation, peptide loading was verified by β2-microglobulin staining. DMSO served as the negative loading control. PEP-1990 could not be loaded onto T2 cells.

### Structural validation of the ParaPredict VR-4 and VR-6 models against the corresponding cryo-EM structures

VR-4 and VR-6 Fab-pHLA complexes were modeled using ParaPredict, our integrated structure-prediction workflow that combines Chai-1 based ab initio modeling with restraints derived from prior structural and experimental (laboratory) data to guide the docking of the Fab antibody onto the pHLA antigen, together with a composite scoring algorithm to rank the resulting antibody-pHLA complex models [18]. To test how the resulting top-ranked ParaPredict models reflect the cryo-EM determined complexes, the top VR-4 model and the top VR-6 model were superposed directly onto the deposited cryo-EM coordinates. Model and cryo-EM chains were matched by sequence identity, and three complementary structural metrics were evaluated: (i) RMSD of the pHLA moiety (MHC heavy chain and peptide), (ii) RMSD of the CDR loops (all CDRs, CDR-H3, CDR-L3) after superposition on the non-CDR Fv framework, and (iii) the residual CDR displacement after superposition on the pHLA reference frame, reflecting the accuracy of the global Fab-antigen docking geometry. In parallel, the calculated (MOE) contact and interaction energies for each ParaPredict model and its cryo-EM counterpart were compared, classifying every contact as ionic hydrogen bond (salt bridge), hydrogen bond, arene contact, or generic (van der Waals-range) contact, and as common to model and cryo-EM structure, model-only, or cryo-EM-only.

For VR-4 (Figure S8A), the pHLA moiety superposed onto the cryo-EM structure with an RMSD of 0.554 Å, confirming accurate modeling of the HLA-A*02:01/MAGE-A4 antigen. CDR loop RMSDs (1.567 Å all CDRs, 1.610 Å CDR-H3, 1.780 Å CDR-L3) indicate good agreement for these conformationally flexible loops, and the CDR displacement after pHLA-frame superposition (1.473 Å) confirms correct overall Fab-pHLA modeling. The contact comparison corroborates this picture quantitatively: of the three antibody-peptide salt bridges present in the cryo-EM structure, two are reproduced by the model with near-identical geometry and energy (CDR-H2 Glu51-Arg6: 2.66 Å /−37.2 kcal/mol in the model vs. 2.71 Å /−34.0 kcal/mol in the cryo-EM structure; Asp99-Arg6: 2.68 Å /−30.5 kcal/mol vs. 2.73 Å /−22.0 kcal/mol), and the single antibody-HLA salt bridge (CDR-L2 Glu55-Arg66) is reproduced almost exactly (2.73 Å /−31.9 kcal/mol vs. 2.75 Å /−37.5 kcal/mol). The third peptide-directed salt bridge, CDR-H1 Lys33-Asp4 (−20.9 kcal/mol in the cryo-EM structure), is absent from the model (closest approach 4.88 Å) despite an accurately placed CDR-H1 backbone (loop RMSD 0.47 Å), pointing to a sidechain rotamer-level rather than a loop-conformation discrepancy. Of the 15 polar/arene contacts identified in the cryo-EM structure, 10 are also present in the model; the remaining differences are limited to weak, peripheral hydrogen bonds (ΔE ¡ 8 kcal/mol). Consistent with this close agreement at the level of individual contacts, the total calculated interaction energies are closely matched between model and cryo-EM structure (antibody-peptide: −82.6 vs. −87.5 kcal/mol; antibody-HLA: −58.5 vs. −52.0 kcal//·/mol; combined: −141.0 vs. −139.5 kcal/mol), indicating that the single missing salt bridge is energetically compensated by small differences elsewhere in the interface rather than reflecting a systematic over- or under-prediction of binding.

For VR-6 (Figure S8B), the pHLA moiety superposed with an RMSD of 0.657 Å. Isolated CDR loop RMSDs were low (0.718 Å all CDRs, 0.629 Å CDR-H3, 1.098 Å CDR-L3), reflecting precise prediction of the individual loop conformations, but the CDR displacement after pHLA-frame superposition was substantially larger (4.883 Å), indicating a modest rigid-body mis-orientation of the Fab relative to the pHLA complex. However, both key salt bridges of the cryo-EM structure are reproduced by the model with near-identical geometry and energy (CDR-H3 Glu99-Arg6: 2.70 Å /−27.2 kcal/mol vs. 2.76 Å /−31.8 kcal/mol; CDR-H2 Arg54-Glu167: 2.73 Å /−38.4 kcal/mol vs. 2.71 Å /−31.7 kcal/mol). Due to this geometry displacement, the model forms an additional, energetically substantial salt bridge that is not found in the cryo-EM structure: CDR-H3 Asp107-Arg66 (α1 helix), contributing −28.4 kcal/mol in the model. This contact co-locates with the CDR-H3/HLA-α1 surface implicated in the pHLA-frame CDR displacement, consistent with a slight global rotation of the Fab that brings an additional heavy-chain acidic residue into proximity with the HLA helix. Five of the polar/arene contacts are common to model and cryo-EM structure; however, the model forms several additional contacts not seen in the cryo-EM structure (predominantly weak antibody-HLA hydrogen bonds, e.g., Thr28-Glu59, Tyr32-Glu59). This more extensive, energetically weighted mismatch in the secondary contact network is directly reflected in the total interaction energies: whereas the antibody-peptide energy is similar between model and cryo-EM structure (−48.6 vs. −52.2 kcal/mol), the antibody-HLA energy is nearly twice as favorable in the model as in the cryo-EM structure (−96.4 vs. −49.7 kcal/mol), driven principally by the non-native Asp107-Arg66 salt bridge and the additional model-only hydrogen bonds. The ParaPredict-derived VR-6 model thus over-predicts the strength of the antibody-HLA interface specifically, providing an energetic signature of the same global docking-orientation error already identified at the structural level.

Taken together, both ParaPredict models reproduce the peptide-centric, TCR-like binding mode with near-quantitative accuracy at the level of the dominant, interface-defining salt bridges: three of the four verified antibody-peptide/antibody-HLA salt bridges are reproduced within 0.06 Å and, for VR-4, within a few kcal/mol of the experimental energy. The residual discrepancies are mechanistically distinct and spatially confined rather than diffuse: for VR-4, a single side-chain-level contact (CDR-H1 Lys33-Asp4) is lost despite correct backbone placement, with no net effect on the total interaction energy; for VR-6, a modest global shift of the Fab orientation produces one additional, energetically substantial non-native contact (Asp107-Arg66) that inflates the predicted antibody-HLA binding energy. This localizes the modeling error to specific, addressable features of the docking geometry rather than a general failure of the restraint-guided prediction.

**Figure S8.**
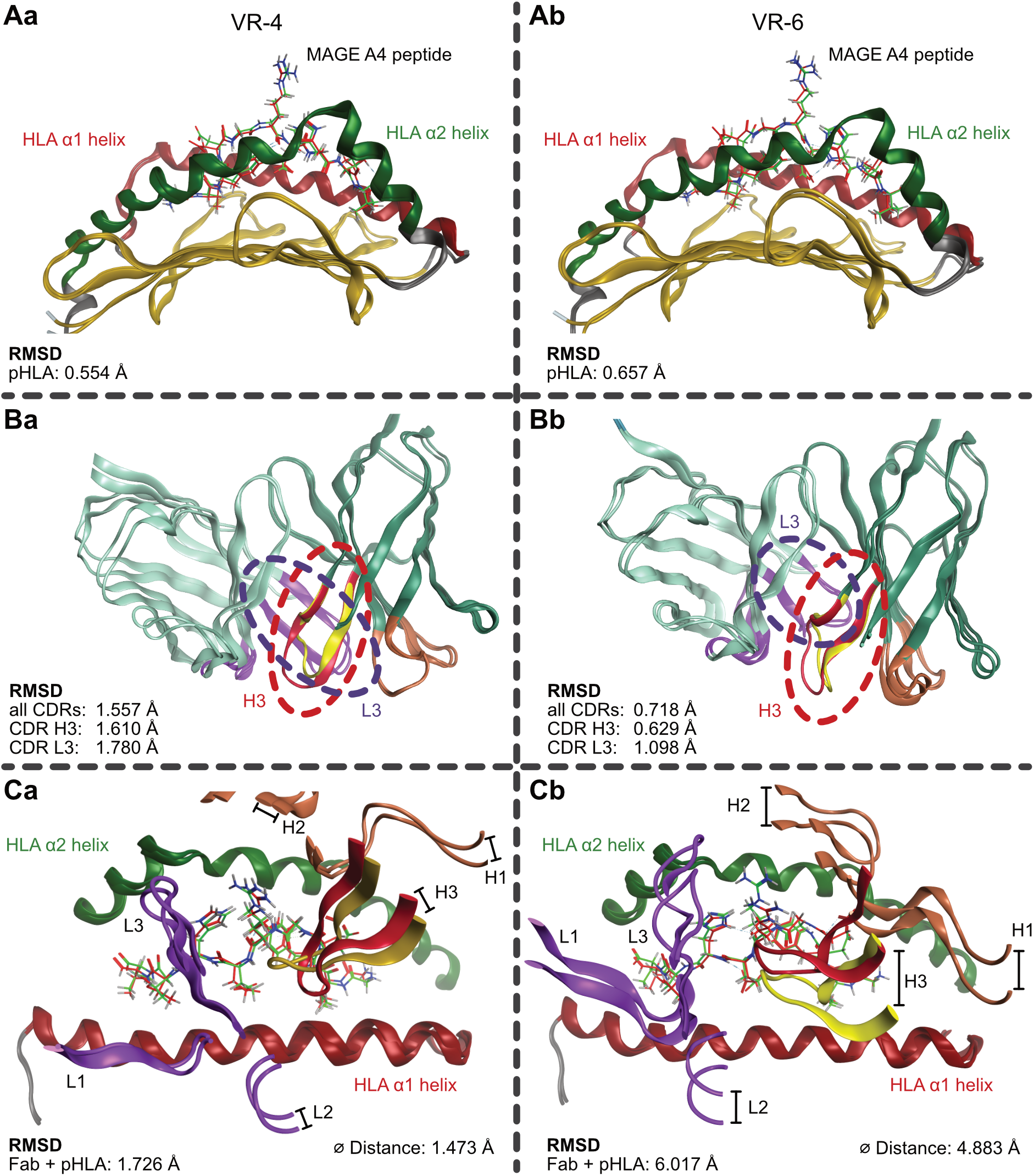
Structural validation of the ParaPredict VR-4 and VR-6 models against the corresponding cryo-EM structures. Top-ranked ParaPredict models for the VR-4 and VR-6 Fab-pHLA complexes were superposed onto the corresponding cryo-EM coordinates and compared using three complementary metrics: RMSD of the pHLA moiety, RMSD of the CDR loops (all CDRs, CDR-H3, CDR-L3) after pairwise superposition of the CDR atoms. **(Aa, Ab)** Cryo-EM structure and model of the pHLA complex (ribbon and sticks, colored by domain: HLA α1 helix, red; α2 helix, green; β-sheet, gold; peptide cryo-EM structure, green; peptide model, red) superposed for VR-4 **(Aa)** and VR-6 **(Ab)**. The pHLA moiety is accurately reproduced in both models (RMSD: VR-4, 0.554 Å; VR-6, 0.657 Å). **(Ba, Bb)** CDR loop superposition, with CDR-H3 (cryo-EM structure, red; model, yellow) and CDR-L3 (purple) highlighted (dashed outlines), for VR-4 **(Ba)** and VR-6 **(Bb)**. CDR loop conformations are well predicted in both models (all CDRs/CDR-H3/CDR-L3 RMSD: VR-4, 1.557/1.610/1.780 Å; VR-6, 0.718/0.629/1.098 Å). **(Ca, Cb)** Full Fab-pHLA complexes superposed, showing the position of the CDR loops (H1-H3, L1-L3) relative to the HLA α1 helix and α2 helix for VR-4 **(Ca)** and VR-6 **(Cb)**. Coloring of the helices, peptides and CDR-loops as in Aa, Ab, Ba, Bb. RMSD of the complex and mean residual displacement (indicated black bar, four distances determined and shown as average distance value) of the Fab quantify global docking accuracy (VR-4: RMSD 1.726 Å, Ø1.473 Å; VR-6: RMSD 6.017 Å, Ø4.883 Å), indicating accurate global docking for VR-4 and a modest rigid-body mis-orientation for VR-6. Hydrogen atoms were added computationally for visualization and hydrogen-bond assignment.

**Figure S9.**
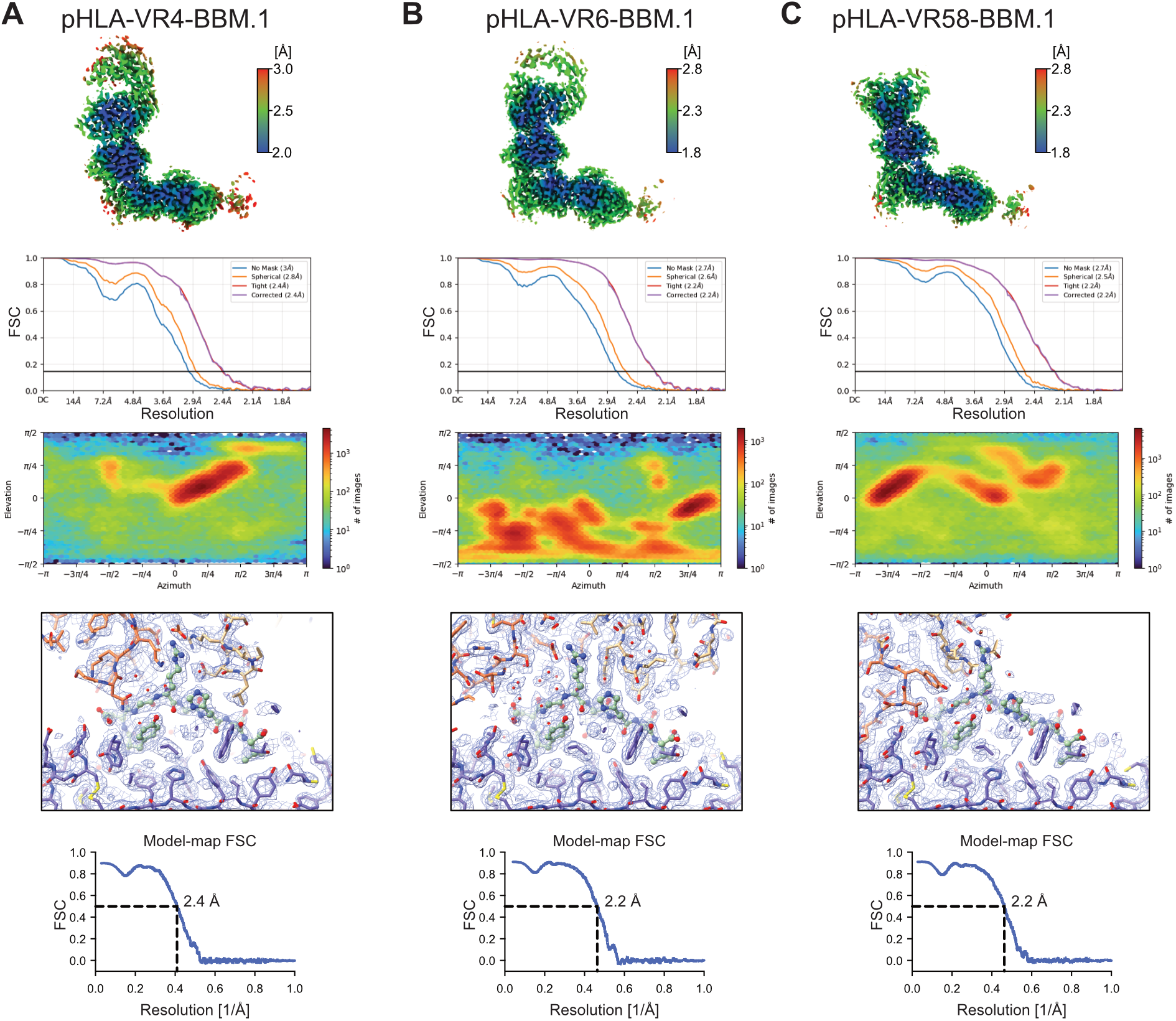
Cryo-EM reconstruction for pHLA-Fab complexes. From top to bottom: local resolution estimation, Fourier shell correlation (FSC) curves, angular distribution of particles, cryo-EM densities around MAGE-A4 peptide, and cryo-EM map-to-model fitting FSCs of the final cryo-EM maps of pHLA in complex with VR-4 **(A)**, VR-6 **(B)**, or VR-58 **(C)**. The local resolution maps are shown as cut-away views through the peptide binding site of HLA. MAGE-A4 peptide, green; HLA, purple; VR Fabs light chain, beige; VR Fabs heavy chain, orange.

**Figure S10.**
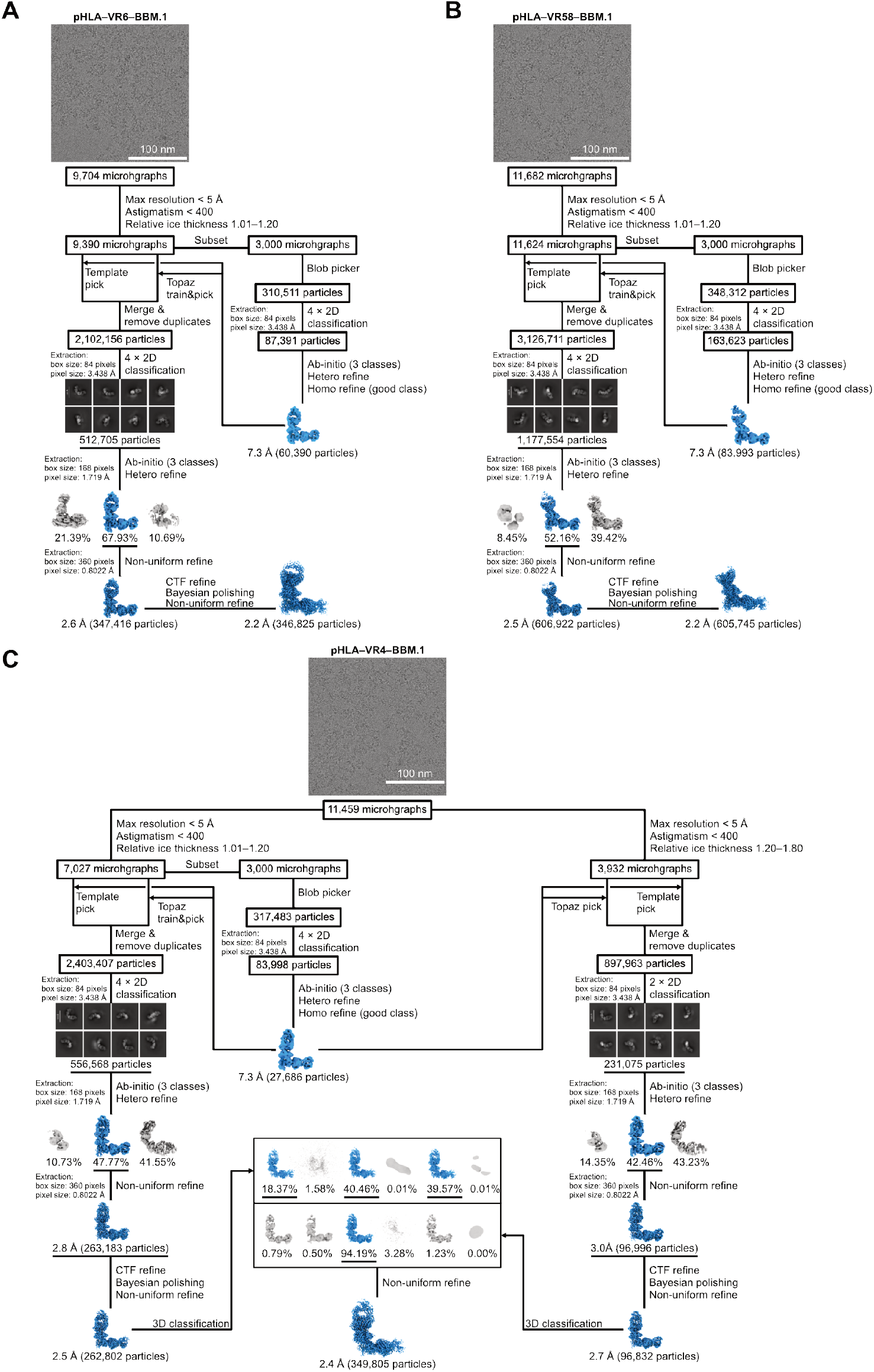
Flow chart single-particle cryo-EM analysis of pHLA-VR Fab complexes. **(A-C)** Data processing pipeline of pHLA in complex with Fab VR-6, VR-58, and VR-4, respectively.

**Table S1.** EpiTox-predicted binder annotation table. (not included in preprint; available with the peer-reviewed publication)

**Table S2.** EpiPredict-predicted binder annotation table. (not included in preprint; available with the peer-reviewed publication)

**Table S3.** Comparative structural parameters of anti-MAGE-A4 TCRm-pHLA complexes.

| Feature | VR-4 | VR-6 | VR-58 |
| --- | --- | --- | --- |
| Ab-contacts to peptide position | P3-P9 | P3-P9 | P4-P6 |
| Total Ab-peptide contacts | 20 | 23 | 13 |
| Ab-peptide H-bonds / salt bridges | 6/3 | 7/1 | 5/1 |
| Dominant Ab-peptide contact | CDR-H2 Glu51-Arg6 | CDR-H3 Glu99-Arg6 | CDR-L2 Asp50-Arg6 |
| Ab-HLA helix contacts | 29 | 35 | 41 |
| Ab-HLA H-bonds / salt bridges | 9/1 | 4/1 | 13/1 |
| Strongest Ab-HLA contact | CDR-L2 Glu55-Arg66 ($\alpha 1$) | CDR-H2 Arg54-Glu167 ($\alpha 2$) | CDR-L1 Arg29-Glu155 ($\alpha 2$) |
| $\alpha 2:\alpha 1$ contact balance | balanced (1.4) | balanced (1.3) | $\alpha 2$-dominant (6) |
| Binding geometry | central / TCR-like | central / TCR-like | shifted toward $\alpha 2$ (F-pocket) |
| Calculated E Ab-pHLA (kcal/mol)* | -140 | -102 | -96 |
| Calculated E Ab-peptide (kcal/mol)* | -88 | -52 | -47 |
| Calculated E Ab-HLA (kcal/mol)* | -52 | -50 | -49 |

**Table S4.**
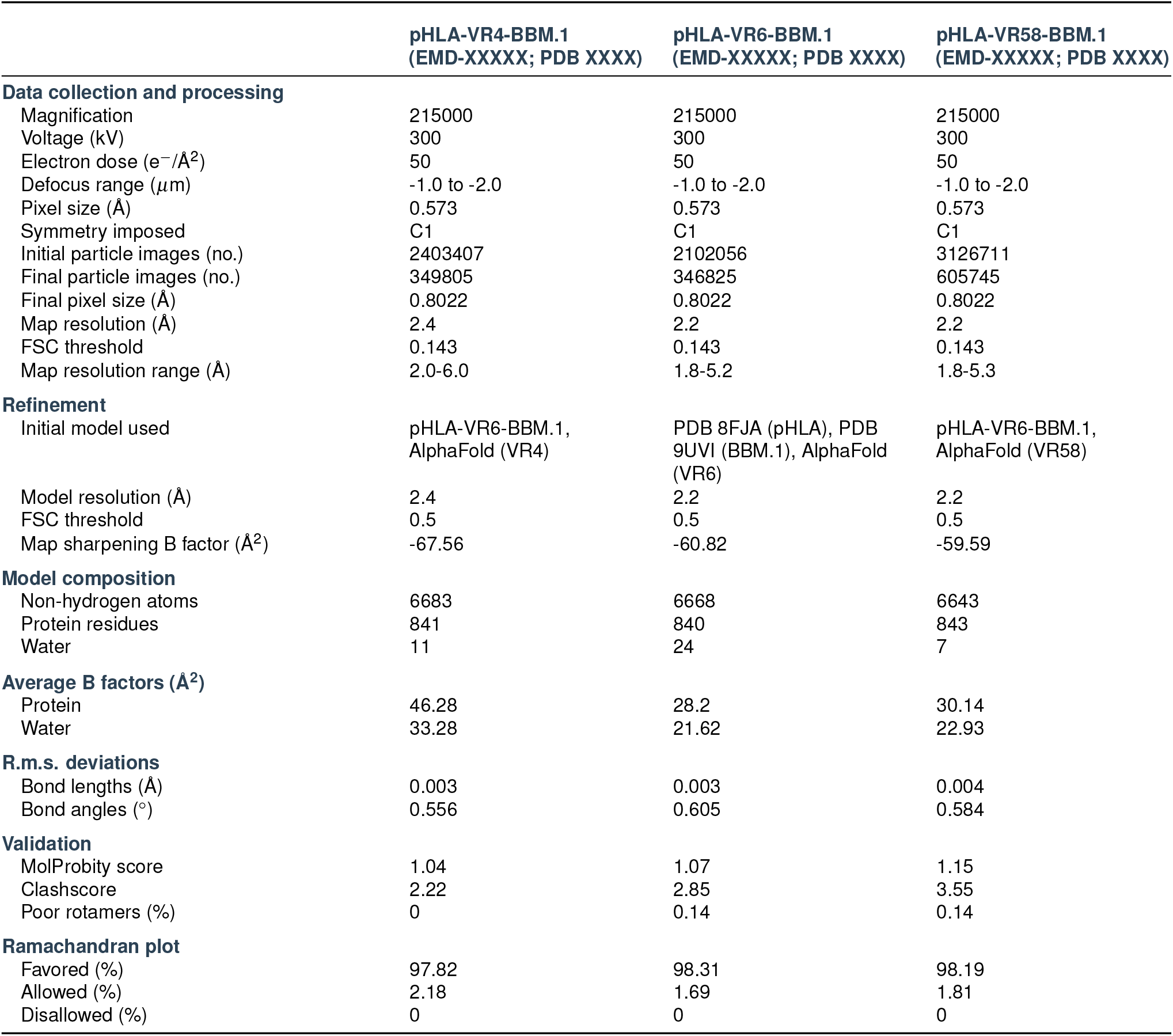
Cryo-EM and model data statistics.

|  | pHLA-VR4-BBM.1<br>(EMD-XXXXX; PDB XXXX) | pHLA-VR6-BBM.1<br>(EMD-XXXXX; PDB XXXX) | pHLA-VR58-BBM.1<br>(EMD-XXXXX; PDB XXXX) |
| --- | --- | --- | --- |
| **Data collection and processing** |  |  |  |
| Magnification | 215000 | 215000 | 215000 |
| Voltage (kV) | 300 | 300 | 300 |
| Electron dose ($e^-/\text{\AA}^2$) | 50 | 50 | 50 |
| Defocus range ($\mu\text{m}$) | -1.0 to -2.0 | -1.0 to -2.0 | -1.0 to -2.0 |
| Pixel size ($\text{\AA}$) | 0.573 | 0.573 | 0.573 |
| Symmetry imposed | C1 | C1 | C1 |
| Initial particle images (no.) | 2403407 | 2102056 | 3126711 |
| Final particle images (no.) | 349805 | 346825 | 605745 |
| Final pixel size ($\text{\AA}$) | 0.8022 | 0.8022 | 0.8022 |
| Map resolution ($\text{\AA}$) | 2.4 | 2.2 | 2.2 |
| FSC threshold | 0.143 | 0.143 | 0.143 |
| Map resolution range ($\text{\AA}$) | 2.0-6.0 | 1.8-5.2 | 1.8-5.3 |
| **Refinement** |  |  |  |
| Initial model used | pHLA-VR6-BBM.1, AlphaFold (VR4) | PDB 8FJA (pHLA), PDB 9UVI (BBM.1), AlphaFold (VR6) | pHLA-VR6-BBM.1, AlphaFold (VR58) |
| Model resolution ($\text{\AA}$) | 2.4 | 2.2 | 2.2 |
| FSC threshold | 0.5 | 0.5 | 0.5 |
| Map sharpening B factor ($\text{\AA}^2$) | -67.56 | -60.82 | -59.59 |
| **Model composition** |  |  |  |
| Non-hydrogen atoms | 6683 | 6668 | 6643 |
| Protein residues | 841 | 840 | 843 |
| Water | 11 | 24 | 7 |
| **Average B factors ($\text{\AA}^2$)** |  |  |  |
| Protein | 46.28 | 28.2 | 30.14 |
| Water | 33.28 | 21.62 | 22.93 |
| **R.m.s. deviations** |  |  |  |
| Bond lengths ($\text{\AA}$) | 0.003 | 0.003 | 0.004 |
| Bond angles ($^\circ$) | 0.556 | 0.605 | 0.584 |
| **Validation** |  |  |  |
| MolProbity score | 1.04 | 1.07 | 1.15 |
| Clashscore | 2.22 | 2.85 | 3.55 |
| Poor rotamers (%) | 0 | 0.14 | 0.14 |
| **Ramachandran plot** |  |  |  |
| Favored (%) | 97.82 | 98.31 | 98.19 |
| Allowed (%) | 2.18 | 1.69 | 1.81 |
| Disallowed (%) | 0 | 0 | 0 |

